# Structure of the thin filament in native skeletal muscles reveals its interaction with nebulin and two distinct conformations of myosin

**DOI:** 10.1101/2021.10.06.463400

**Authors:** Zhexin Wang, Michael Grange, Sabrina Pospich, Thorsten Wagner, Ay Lin Kho, Mathias Gautel, Stefan Raunser

**Affiliations:** Department of Structural Biochemistry, Max Planck Institute of Molecular Physiology, Otto-Hahn-Strasse 11, 44227, Dortmund, Germany; Randall Centre for Cell and Molecular Biophysics, School of Basic and Medical Biosciences, Kings College London BHF Centre of Research Excellence, New Hunt’s House, Guy’s Campus, London SE1 1UL, UK; Structural Biology, The Rosalind Franklin Institute, Harwell Science and Innovation Campus, Didcot OX11 0FA, UK

**Author notes:** The authors contributed equally (Zhexin Wang, Michael Grange).

## Abstract

Nebulin is a major structural protein of skeletal sarcomeres and is essential for proper assembly and contraction of skeletal muscle^1^. It stabilises and regulates the length of thin filaments,^2^ but the structural mechanism remains nebulous. Using electron cryotomography and sub-tomogram averaging, we present the first structure of native nebulin bound to thin filaments within the A-band and I-band of intact sarcomeres. This *in-situ* reconstruction reveals unprecedented detail of interaction at pseudo-atomic resolution between nebulin and actin, providing the basis for understanding the structural and regulatory roles of nebulin. The position of nebulin on the thin filament indicates that there is no contact to tropomyosin or myosin, but an unexpected interaction with a troponin-T linker, possibly through two binding motifs on nebulin. In addition, our structure of myosin bound to the thin filaments reveals different conformations of the neck domain, both within the same sarcomere and when compared to purified structures, highlighting an inherent structural variability in muscle. We provide a complete description of cross-bridge formation on fully native, nebulin-containing thin filaments at near-atomic scale. Our structures establish the molecular basis for the role of nebulin as a thin filament “molecular ruler” and the impact of nemaline myopathies mutations that will aid future development of therapies.

## Main text

In skeletal sarcomeres, thin and thick filaments lengths are regulated by two elongated giant proteins, nebulin and titin, respectively^3,4^. A single nebulin molecule (molecular weight over 700 kDa) binds along the entire thin filament from the barbed end in the Z-disc to the pointed end near the M-band^5,6^, maintaining the stability of thin filaments^2^. Mutations in its encoding gene, NEB, are a major cause of a class of skeletal muscle disorders termed nemaline myopathies that present with a range of pathological symptoms such as hypotonia, muscle weakness and, in some cases, respiratory failure leading to death^7–9^. Despite the critical role of nebulin in skeletal muscle, nebulin is only minimally expressed in cardiac muscle^10^, where instead nebulette, a short homolog of nebulin, is present but only close to the Z-disc. The absence of nebulin results in a broader range of thin filament length^11^ in cardiomyocytes that possibly enables greater tunability of activation^12^.

Nebulin primarily consists of 22-28 tandem super repeats. Each super repeat consists of 7 simple repeats, each made of 31-38 amino acid residues, featuring a conserved sequence motif SDxxYK^13,14^. The N- and C-termini of nebulin associate with the capping proteins on the two ends of the thin filaments, tropomodulin (towards the M-band)^15^ and CapZ (at the Z-disc)^16^, respectively. Therefore, nebulin is hypothesised to regulate thin filament length as a “molecular ruler”, albeit with the exact mechanism remaining unknown^2,17–20^. Indeed, genetic ablation of nebulin in mice is lethal and results in sarcomeres with loss of their length-regulation^3,21^.

It has been suggested that, based on the modular sequence of nebulin, each simple repeat would bind to one actin subunit and every seventh repeat, i.e. a super repeat, would interact with the tropomyosin-troponin regulatory complex^13^. However, there is so far no structural detail of these interactions or even of native nebulin itself. Therefore, it remains elusive how nebulin stabilises or even regulates thin filaments. Unfortunately, the enormous size of nebulin combined with its elongated and flexible nature has prevented the use of *in vitro* reconstituted systems of nebulin and thin filaments that would resemble the native state in a sarcomere. Recombinant nebulin fragments, while binding to F-actin, were found to bundle actin filaments^22^, rendering them useless for a reconstitution approach for electron-microscopical structural biology.

In this study, we imaged nebulin directly inside mature mouse skeletal sarcomeres from isolated myofibrils using cryo-focused-ion-beam milling (cryo-FIB) and electron cryotomography (cryo-ET). We present the structure of nebulin bound to the actin filament at a resolution of 4.5 Å as well as a structure of nebulin-containing actomyosin at 6.6 Å. The structures reveal the native conformation of nebulin inside the sarcomere and allow us to describe the molecular interactions underpinning its organisation. Through the comparison of these structures to cardiac thin filament structures, our *in-situ* reconstructions provide us with new insights into the interactions between nebulin and troponin, the arrangement of myosin double-heads, and how the thin filament accommodates nebulin.

### *In situ* position of nebulin on thin filaments

We determined the structure of the core of the thin filament from intact myofibrils isolated from mouse psoas muscle to 4.5 Å resolution and with actomyosin resolved to 6.6 Å resolution (Extended Data Figs. 1,2). This is, to our knowledge, the highest resolution structure of a filament determined *in-situ* and the only pseudo-atomic structure determined from tissues after cryo-FIB. In the core of the thin filament, two extra continuous densities were visible alongside the actin filament (Fig. 1a-c). The elongated structure predicted for nebulin^23^ suggested that this density might be natively organised nebulin bound to the thin filament. To verify this putative identification, we determined the *in situ* actomyosin structure in the A-band from cardiac muscle (Extended Data Fig. 3a,b). It has been previously shown that nebulin is barely expressed only in small subpopulations of myofibrils in cardiac muscle, while nebulette is only located close to Z-disc^10^. The averaged reconstruction of the cardiac thin filament, determined to an overall resolution of 7.7 Å with the core of thin filament resolved to 6.3 Å, depicts similar organisations of actin, myosin and tropomyosin. Importantly, the extra density observed in skeletal actomyosin is missing (Fig. 1e), confirming that this density corresponds to averaged segments of nebulin.

**Fig. 1.**
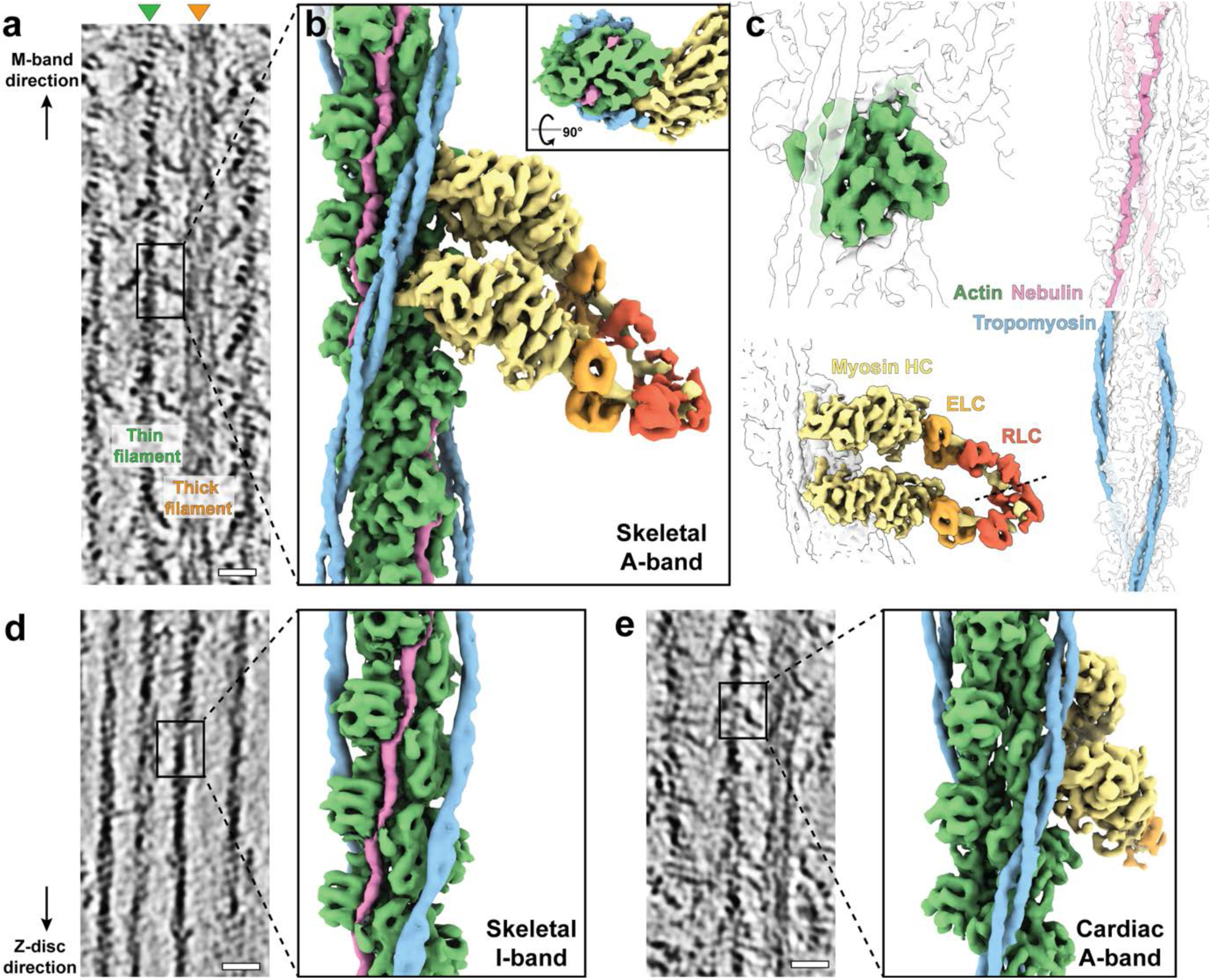
Thin filament structures in striated muscle sarcomeres. **a**. Tomographic slice of skeletal sarcomere A-band depicting adjacent thin and thick filaments. **b**. Actomyosin structure from the skeletal sarcomere A-band consisting of actin (green), myosin (heavy chain (HC): yellow, essential light chain (ELC): orange, regulatory light chain (RLC): red), tropomyosin (blue) and nebulin (magenta). Myosin is a composite map including light chains from different averaged structures (see Extended Data Figs. 1,9). Cross-section view of the structure is shown in the inset. **c**. Different components of a thin filament and their position highlighted within the structure. The dotted line highlights the interface between the two RLCs of the upper and lower myosin head. **d**. Tomographic slice of a skeletal sarcomere I-band and structure of the thin filament (inset). **e**. Tomographic slice of a cardiac sarcomere A-band and structure of actomyosin, including a pair of myosin double-heads. All tomographic slices are 7 nm thick. Scale bars: 20 nm

Nebulin lies in the grooves between the two strands of the actin filament, following their helical turn (Fig. 1a,2b). Interestingly, nebulin occupies a site that is known to be bound by actin-stabilising compounds such as phalloidin and jasplakinolide^24^ (Extended Data Fig. 4), which explains why excessive phalloidin can unzip nebulin from thin filaments^25^ and may also explain a similar mechanism of F-actin stabilisation. A single actin filament is decorated by two nebulin molecules on the opposite sides (Fig. 1b). In order to ascertain the molecular organisation of nebulin in different regions of a sarcomere, we also determined the structure of the thin filament in the skeletal muscle I-band to a resolution of 7.4 Å (Extended Data Fig. 3c,d). Nebulin appears in the I-band at the same position on the thin filament as is observed within the A-band (Fig. 1d), indicating that nebulin spans most of the thin filament^19,26^. This supports the general concept that nebulin acts as a “molecular ruler”^17,27^ and suggests that nebulin maintains a structural role within the sarcomere. Significantly, the position of nebulin bound to actin from native skeletal muscle is different from the three putative sites previously proposed on the outer surface of the actin filament based on reconstituted actin-nebulin fragment complexes^28^. The observed differences could represent the limitations of the use of *in-vitro* fragments of nebulin or suggest that there are different interaction patterns during sarcomerogenesis.

The position of nebulin implies that it does not interact with tropomyosin or myosin (Fig. 1b). The physical separation by the subdomain 3 and 4 (SD3, 4) of adjacent actin monomers ensures that nebulin is never in close contact with tropomyosin, regardless of the tropomyosin state at different Ca^2+^ concentrations^29^ (Extended Data Fig. 5), contradictory to previous results from *in-vitro* experiments^30^. The discrepancy between our *in situ* structures and *in vitro* assays again demonstrates that nebulin may have different properties when purified, compared to its native state in a sarcomere. Purified large fragments of nebulin are extremely insoluble when expressed^22,31,32^. Both rotary shadowed images of nebulin^32^ and the structure of nebulin predicted by the machine-learning-based software, AlphaFold^33^, suggest compact globular structures. These visualisations strongly deviate from the elongated shape of nebulin when bound to actin filaments. Therefore, our approach of investigating nebulin inside sarcomeres provides in situ structural information about nebulin interactions with the thin filament that are not accessible by sequence-based structure prediction programs or from isolated proteins. These discrepancies also suggest that during sarcomerogenesis, nebulin integration into the thin filament is likely to require cellular cofactors preventing the formation of aggregates or large globular structures, which remain to be identified.

### Nebulin structure and localisation of residues

While nebulin consists of repetitive simple repeats, each simple repeat has different sequences, with a few conserved charged residues and a putative actin-binding SDxxYK motif (Fig. 2a,g, Extended Data Fig. 6a). Due to the nature of sub-tomogram averaging, the obtained EM density map of nebulin is an averaged density of all repeats in the A-band. Taking advantage of the 4.5 Å map, where bulky side chains are typically resolved (Extended Data Fig. 2), we were able to build an atomic model for actin and refine a poly-alanine nebulin model into its density (Extended Data Table 1). Using a published convention^13^, we defined the start of a simple repeat at two residues preceding a conserved aspartic acid, resulting in the SDxxYK motif residing at position 18-23.

**Fig. 2.**
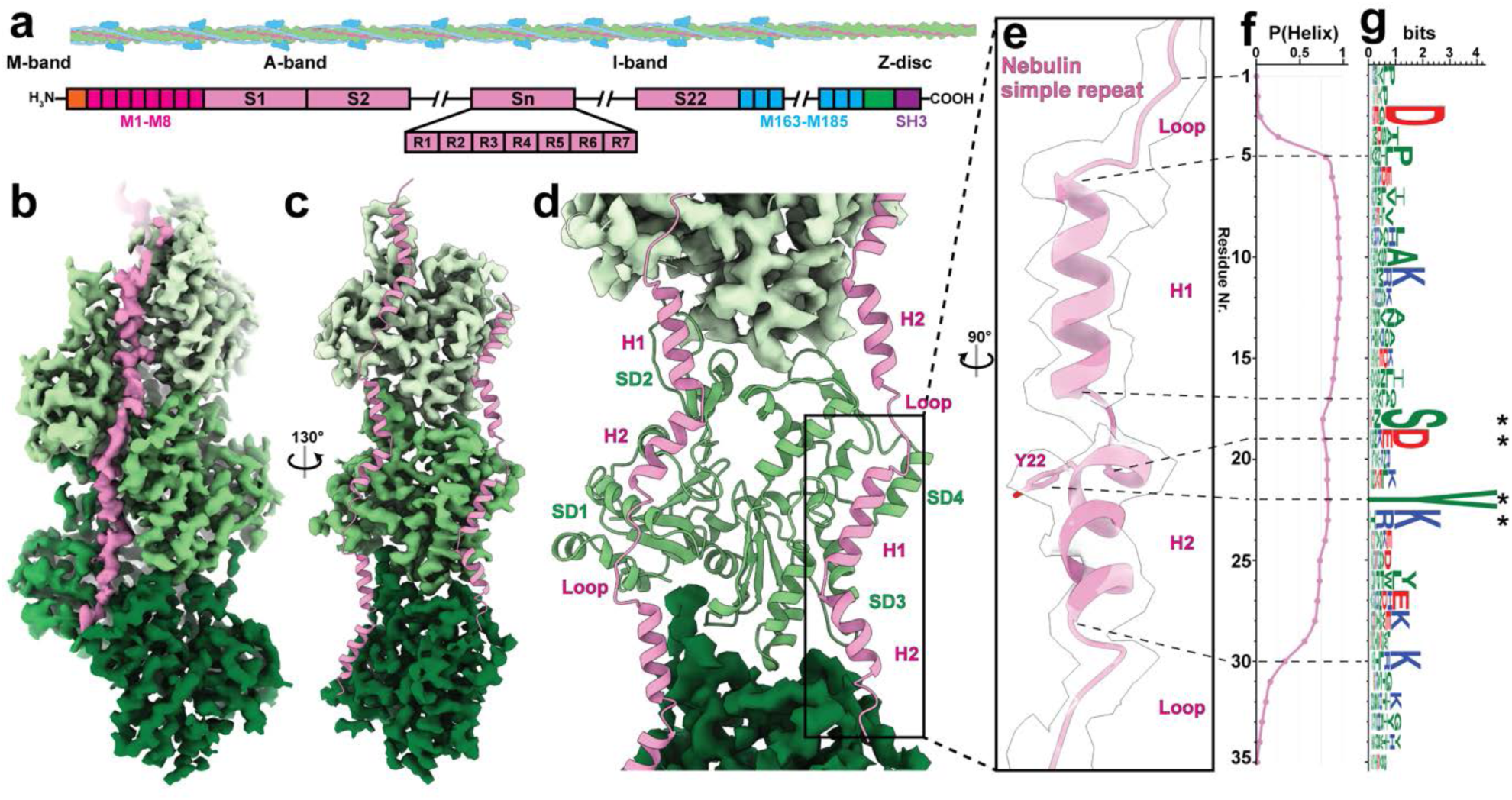
Nebulin structure and its binding to the actin filament. **a**. Schematic drawing of the nebulin-bound thin filament and modular organisation of the primary sequence of nebulin demonstrating its super repeats. **b**. Sub-tomogram averaged structure of the actin filament in complex with nebulin (magenta) at a resolution of 4.5 Å. Different actin subunits are coloured in different shades of green with darker green towards the barbed end. **c**. Rotated view of (**b**) highlighting both nebulin molecules (shown as structural models of three and two simple repeats) on the actin filament. Only one strand of the actin filament is shown. **d**. Structural model of one actin subunit and two nebulin molecules. One nebulin binds along actin subdomain 1 and 2 (SD1 and SD2) while the other binds along actin subdomain 3 and 4 (SD3 and SD4). The cryo-EM map of the neighbouring actin subunits is shown. **e**. Zoom-in view of one nebulin simple repeat. The side chain of residue Y22 is highlighted. **F**. Averaged predicted score for an α-helix at each residue position of a simple repeat. **g**. Graphical representation of sequence alignment of all simple repeats (M1-M163). A larger amino acid symbol corresponds to a greater occurrence at a certain position. Positive, negative and neutral residues are coloured in blue, red and green, respectively. Dotted lines map the sequence to the structural model in (**e**) and (**f**). Asterisks mark the conserved SDxxYK motif.

The model of nebulin consists of a repetitive structure of two α-helices (H1 and H2), with a short kink of 46° in between, followed by a loop region spanning around SD1 of actin (Fig. 2b-d). As validation, and in order to map the sequence to our structural model, we predicted the average secondary structure to highlight structured and unstructured regions from the sequences of nebulin simple repeats (Extended Data Fig. 6b, Methods). The prediction implied each nebulin simple repeat should form a long helix, with a drop in probability in the middle of this helix (Fig. 2f). By matching the predicted start of the helix in the sequence with the start of H1 in the structure, the predicted end of the helix matched the end of H2 and the dip in probability matched the position of the kink in our model (Fig. 2e, f). Based on this registry, a noticeable bulky side chain density aligned with position 22, corresponding to a fully conserved tyrosine residue. We attributed this density as the phenyl group of this tyrosine (Fig. 2e, Extended Data Fig. 2d). This observation further validates the sequence-structure mapping. As such, H1 starts at position 5, which is often occupied by a proline (Fig. 2e-g). The SDxxYK motif, where the exon boundaries are, is located at the beginning of H2, among which the serine is positioned at the kink between H1 and H2. This registry allowed us to assign the location of other conserved residues, and further investigate their roles in the interactions between nebulin and the thin filament..

While most nebulin simple repeats contain 35 amino acids (as is modelled above), some repeats can be as short as 31 aa or as long as 38 aa (Extended Data Fig. 6c). The predicted secondary structure implies that in the shorter repeats, the helix ends earlier than in an average-length repeat and in the longer repeats, the loop is longer (Extended Data Fig. 6d). Although we did not observe separate classes within our cryo-ET data for these repeats due to their low abundancy, it is noticeable that the density corresponding to H2 and the first half of the loop has lower occupancy compared to H1 (Extended Data Fig. 6e). This suggests that in the shorter nebulin repeats, part of H2 is extruded into the loop along segments of actin to compensate for fewer amino acids while in the longer nebulin repeats, the extra amino acids reside flexibly in the loop (Extended Data Fig. 6f). This ensures that in all regions of the sarcomere, nebulin repeats have the same physical length to span an actin subunit to maintain a 1:1 binding stoichiometry and fulfil its role as a “molecular ruler”.

### Interactions between nebulin and the thin filament

Based on our model, we were able to show that the interactions between actin and nebulin are mediated by residues throughout one nebulin simple repeat and three adjacent actin subunits (Fig. 3a). In the SDxxYK motif, Y22 forms a potential cation-pi interaction with K68 on SD1 of one actin subunit (N) (Fig. 3c). S18 likely forms a hydrogen bond with E276 on SD3 of the laterally adjacent actin subunit on the other strand (N+1). D19 and K23 interacts with residues on SD1 and SD2 of actin subunit N through electrostatic attractions. In addition, other highly conserved charged residues outside the SDxxYK motif are also involved in the interactions between actin and nebulin. D3, K11 and K30 can form electrostatic interactions with SD1 of actin subunit N+2, SD4 of actin subunit N+1 and SD1 of actin subunit N, respectively (Fig. 3c). Therefore, the entire nebulin molecule can interact with every actin subunit on the thin filament (Fig. 3a), which prevents them from depolymerisation and confers rigidity and mechanical stability to the thin filament.

**Fig. 3.**
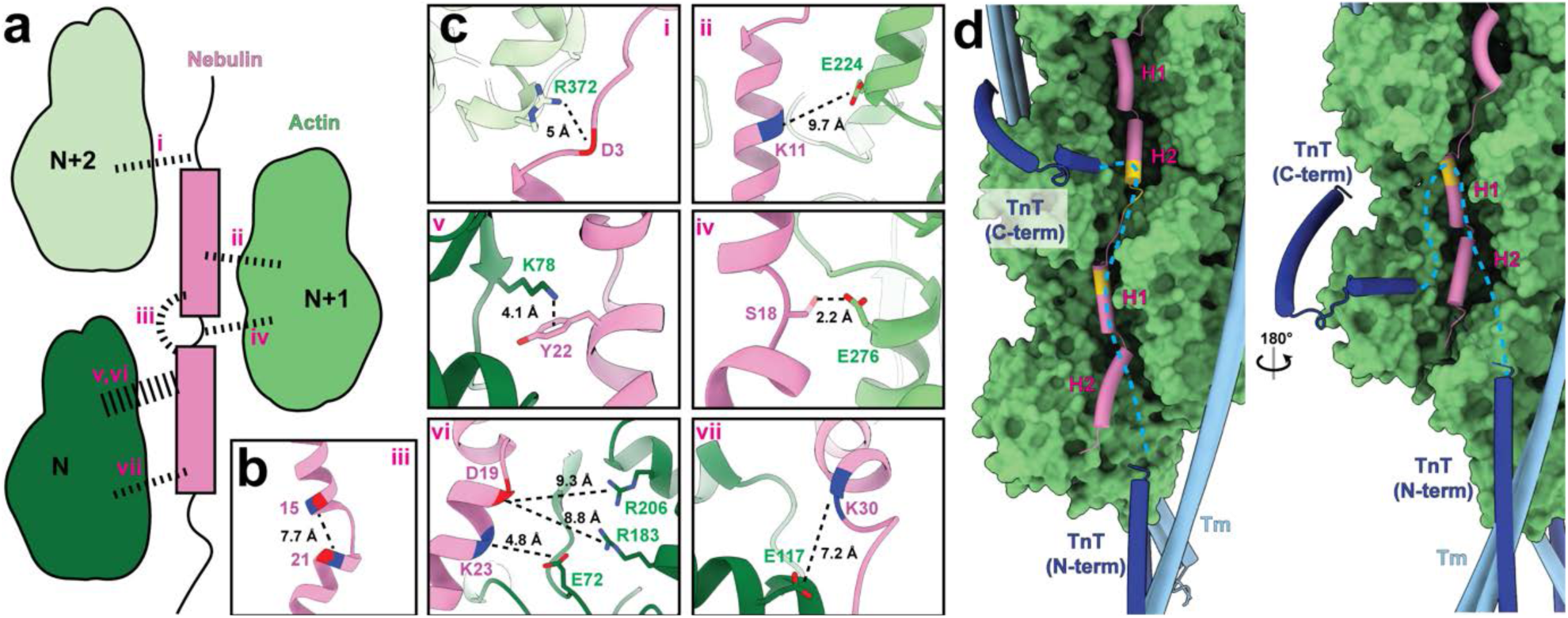
Interactions between nebulin and actin. **a**. Schematic depiction of interactions between nebulin (magenta) and three adjacent actin (green) subunits. Interactions are marked as dotted lines. **b**. Intra-nebulin interactions (iii in (**a**)) between residues with complementary charges at position 15 and 21. **c**. Details of interactions i-vii in (**a**). Distances were measured between actin residues and the Cβ of the poly-alanine model of nebulin where side chain is not resolved. A potential side chain conformation of S18 is shown for visualisation although it was not determined from the map. **e**. Interactions between nebulin and troponin T (TnT). Two different TnT models (dark blue) that bind to opposite sides of the actin filament (PDB: 6KN8) are shown. Possible shapes of the TnT linker are marked as cyan dotted lines based on weak EM densities (EMD-0729). Potential TnT binding sites on nebulin are highlighted in yellow. Tropomyosin (Tm) is shown in light blue.

Interestingly, an intra-molecule interaction occurs between position 15 and 21 on nebulin at the position of the kink between H1 and H2 (Fig. 3a,b). Although both positions can accommodate either positively or negatively charged residues, they appear to be often complementary to each other among all repeat sequences (Extended Data Fig. 7a-c). Their interaction is also supported by weak side-chain densities in our averaged reconstruction (Extended Data Fig. 7d). This intra-molecular interaction stabilises the kink conformation of the two helices, which is necessary for positioning charged residues near actin.

Nebulin simple repeats share a higher sequence similarity with the repeats that are six repeats apart, forming a seven-repeat super repeat pattern (Fig. 2a). This modular structure suggests an interaction with the troponin-tropomyosin regulatory complex, which also has a 1:7 stoichiometry ratio to actin. The physical separation by actin between nebulin and tropomyosin has ruled out their interactions. The core of troponin, including troponin C (TnC), troponin I (TnI) and the majority of troponin T (TnT) are also located away from nebulin (Extended Data Fig. 8a, c). On the other hand, a linker region in TnT between R134 and R179 is likely to be the binding partner of nebulin (Fig. 3d). It was hypothesised to cross the groove between two actin stands^34^. The structures of troponin with actin, reported from cardiac thin filament^35,36^, also show that this TnT linker is localised close to the region where nebulin resides in our structure, although a structural model for the linker could not be determined due to its flexibility. Superimposing the densities for TnT with our structural model of actin and nebulin suggests the location of two contact sites between TnT and nebulin (Extended Data Fig. 8a-d), with one site at the end of H2. This site corresponds to the position of a WLKGIGW motif at the end of repeat 3 in nebulin, which has previously been proposed to be the tropomyosin-troponin binding motif^13^ (Fig. 3d, Extended Data Fig. 8b,f). The other site is located downstream of the first site at the start of H1, indicating another novel potential troponin-binding motif, ExxK, at the beginning of repeat 4 (Fig. 3d, Extended Data Fig. 8b,d,f). The TnT linker region contains a hydrophobic C-terminus and a highly charged N-terminus matching the orientation of the two binding sites, suggesting the WLKGIGW motif and ExxK motif interact with TnT through hydrophobic and electrostatic interactions, respectively (Extended Data Fig. 8g).

In addition to the structural role of nebulin, it has also been shown that nebulin can increase thin filament activation, promote myosin binding and thus improve the efficiency of contraction^37–39^. Based on our observation that nebulin does not interact with myosin or tropomyosin, the regulatory role of nebulin is likely to be a downstream effect of the interaction with the TnT linker. This linker, despite being outside of the core of the troponin complex and flexible in cardiac muscle, is thought to propagate the calcium signal from the core of the troponin complex to the N-terminus of TnT, which binds to tropomyosin^40^. In skeletal muscle, the interaction between nebulin and the TnT linker may rigidify the linker and thus help to maintain efficient calcium regulation and the subsequent binding of myosin.

The molecular detail of the actin-nebulin interaction enables the understanding of the mechanism underlying the pathogenicity caused by recessive mutations in the NEB gene, which is the major cause of nemaline myopathies^8^. Nemaline myopathy mutations are usually compound heterozygous, with one truncating and one missense variant^8^. Two missense NEB mutations, Ser6366Ile and Thr7382Pro, were identified as founder mutations in the Finnish population^41^. The locations of the two sites on a simple repeat correspond to Ser18 and Thr14 (Extended Data Fig. 7e). A mutation of Ser18 into a hydrophobic isoleucine would disrupt its potential hydrogen bond with actin (Fig. 3c, Extended Data Fig. 7f). A mutation of Thr14 to proline, despite not being at a conserved residue position, can lead to the disruption of H1 helical secondary structure and thus alter the local conformation of nebulin and interfere with actin binding (Extended Data Fig. 7f).

Our structural model of actin and nebulin not only explains how nebulin stabilises thin filaments and functions as a “molecular ruler”, but also maps residues crucial in maintaining these interactions. Our approach determined structures across several tissue types and regions, enabling a comparative analysis of nebulin in its native context. When also applied to clinical genetics, this information should help to determine additional pathogenic interfaces of nebulin variants. This is especially crucial when considering missense variants, where the pathogenicity is often difficult to determine^41^ and will thus aid the early diagnosis of nemaline myopathies and genetic counselling of variant carriers.

### Basis of variability within the myosin double-head

In the rigor state of the sarcomere, two myosin heads from a single myosin molecule are bound to the thin filament in most cross-bridges^42^. Understanding the organisation of the myosin heavy chain (HC), essential light chain (ELC) and regulatory light chain (RLC) in an intact double-head is crucial to explain how cross-bridges can maintain their interaction with the thin filament in a native sarcomere in response to external stresses.

Our previous reconstruction of a myosin double-head was limited to a resolution of 15 Å, which prevented us from determining the proper conformation of the lever arm, ELC and RLC^42^. Now, having a better-resolved structure available (∼9 Å in the neck domain), we were able to accurately fit the lever arms and light chains based on their secondary structure elements (Extended Data Fig. 9d-f), completing the model of the entire myosin double-head including HC, ELC and RLC (Fig. 4a, Extended Data Fig. 9). Interestingly, the angles of the kinks in the lever arm helix are different in the upper and the lower head (Fig. 4a). Especially the kink between the two RLC lobes differs considerably in the two heads, resulting in the clamp-like arrangement of the neck domains (Fig. 4b). The RLC-RLC interface resembles that of the RLCs of the free and blocked head in an interacting-head-motif (IHM) of an inactive myosin^43,44^, but with a rotation of ∼20° (Fig. 4c). Although the motor domains are similarly arranged in the cardiac muscle (Extended Data Fig. 10a), our 12 Å reconstruction of the neck domain clearly demonstrates that the interface between the upper and lower head RLCs is different compared to the skeletal counterpart, resulting in a subtle difference in the arrangement of the two neck domains (Extended Data Fig. 10b). In conclusion, our structures of myosin in the ON state in skeletal and cardiac muscles and previous structures of myosin in the OFF state^43,44^ imply natural variabilities within RLCs and at the RLC-RLC interface that allow a dynamic cooperation between the two myosin heads.

**Fig. 4.**
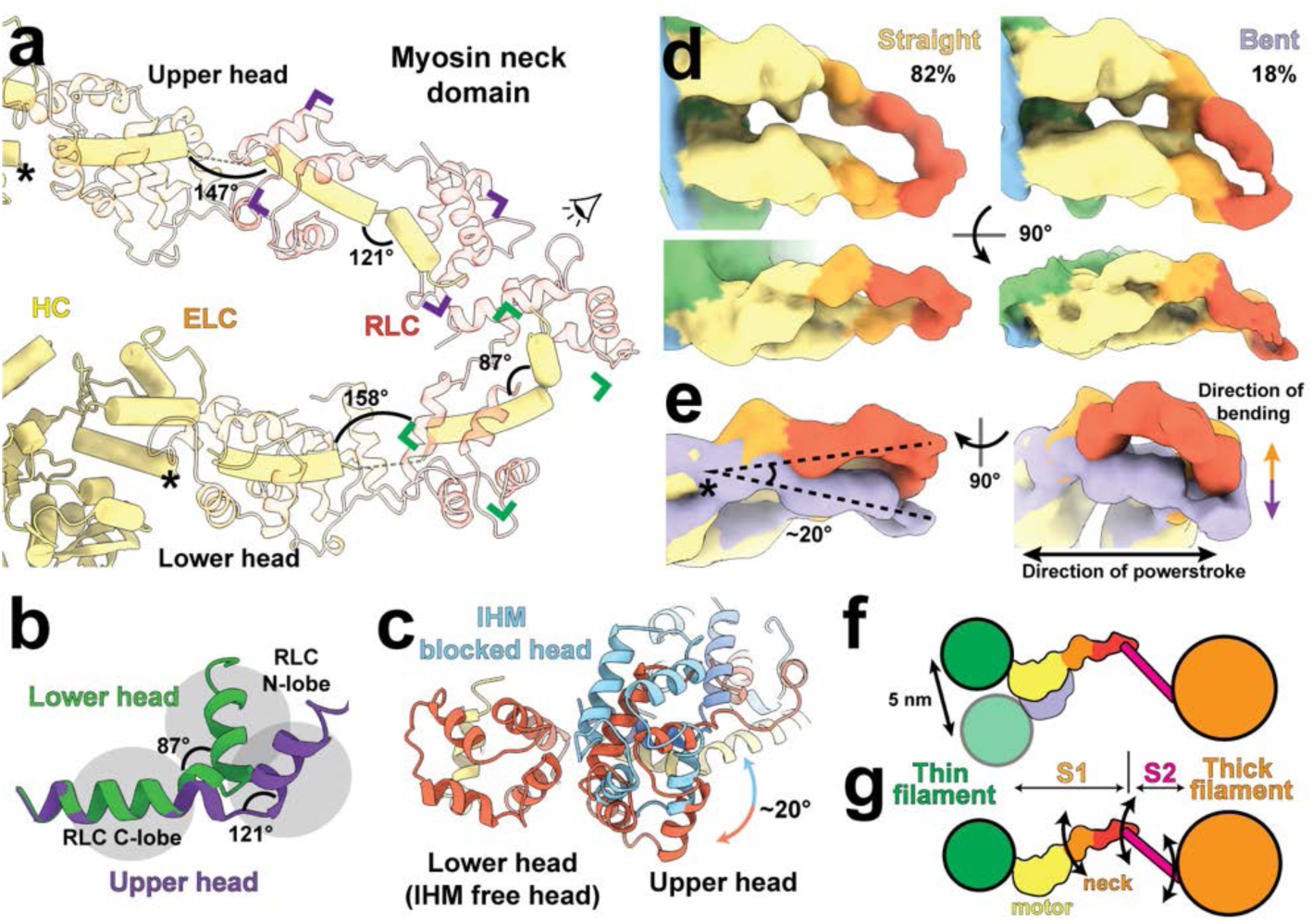
Structural variability within the *in situ* myosin double-head in skeletal muscle. **a**. The lever arms of the upper and lower myosin heads form kinked helices (yellow). The angles at the kinks are different in the upper and the lower head. ELCs and RLCs are shown as transparent models. **b**. Different conformations of the lever arm at the RLC-binding regions of the upper head (purple) and the lower head (green). **c**. View from the eye symbol in (**a**) showing the interface between the RLCs from the upper and lower head (red for RLC, yellow for lever arm helices) compared to the interface of the blocked head (aligned to the lower head) and the free head in the IHM (blue for RLC, dark blue for lever arm helices). **d**. Two different conformations, straight and bent forms, of myosin double heads determined by 3D classification. HC, ELC and RLC regions are coloured in yellow, orange and red. **e**. Comparison between the straight (orange) and bent (purple) double-head conformation. The origin of bending is marked by an asterisk, also in (**a**). **f**. Schematic drawing describing the increased range of thin filament positions that can be bound by myosin heads due to the bending of double-head. **g**. Schematic drawing depicting the three flexible junctions in a myosin head.

Interestingly, 18% of the skeletal double-heads had a different conformation in which both neck domains are bent by ∼20° perpendicular to the direction of the myosin power stroke (Fig 4d,e, Extended Data Fig. 11a). This different structural arrangement increases the range within which myosin can bind to the thin filament by ∼ 5 nm without interfering with force transmission during the power stroke (Fig. 4f). Therefore, the bending contributes additional adaptability on top of what is provided by the flexibility of the S2 domain for cross-bridge formation between actin and myosin filaments (Fig. 4g). The myosin “arm” can therefore hold on tightly to a thin filament, but at the same time have enough freedom to cooperate the mismatch between helical pitches of thick and thin filaments and account for local deformation of the sarcomere. In this line, the double-heads with bent neck domains are randomly distributed in the A-band of the sarcomere (Extended Data Fig. 11b-d), ensuring efficient binding of myosin during contraction.

## Conclusions

Our structural reconstruction of nebulin within a native skeletal sarcomere provides the basis of interaction between nebulin and thin filaments. It reveals the mechanism underlying the stabilising role nebulin plays in the thin filament and its role in regulating myosin-binding through its interaction with TnT. Our approach using cryo-FIB milling and cryo-ET, as the first high-resolution structural study within an isolated tissue, also highlights different conformations of myosin and illustrates similarities and differences from *in vitro* structures. In the context of the sarcomere where several flexible proteins, such as titin and myosin-binding protein-C, are present and still lack structural visualisation, our approach is a general tool for structural analysis where other methods are limited. Determining the structure of these key players in the context of native sarcomeres will enable better modelling of skeletal muscle in the future, directly impacting also the understanding of disease. The structure of nebulin presented here is one such case, where the molecular interactions described might help to establish a foundation for future developments of the treatment of nemaline myopathies. Fundamentally, our structure represents a leap in capability for structural biology. The ability to perform atomic-level structural analyses within native tissue will be at the forefront of molecular diagnostics in the years to come.

## Ethics declarations

Animals were sacrificed in a schedule-1 procedure by cervical dislocation following licensed procedures approved by King’s College London ethics committee and the Home Office UK.

## Acknowledgements

This work was supported by funds from the Max Planck Society (to S.R.), the Wellcome Trust (Collaborative Award in Sciences 201543/Z/16/Z to S.R. and M. Gautel), the European Research Council under the European Union’s Horizon 2020 Programme (ERC-2019-SyG, grant no. 856118 to S.R. and M. Gautel) and the Medical Research Council (MR/R003106/1 to M. Gautel and A.L.K.). M. Grange was supported by an EMBO Long-Term Fellowship. M. Gautel holds the BHF Chair of Molecular Cardiology. We thank S. Tacke for hardware optimisation for cryo-FIB. We are grateful to O. Hofnagel and D. Prumbaum for EM support and B. Brandmeier for technical support.

## Author contributions

S.R. designed and supervised the project. A.L.K. and M. Gautel developed methods and isolated mouse myofibrils. Z.W. performed cryo-FIB and collected cryo-ET data. M. Grange optimised cryo-ET data acquisition. Z.W. and M. Grange performed sub-tomogram averaging.

T.W. implemented automatic filament picking and tools for data conversion and statistical analysis. S.P. built atomic models for actin, nebulin and myosin motor domain. Z.W. performed rigid body fitting of myosin neck domain models. Z.W. prepared figures with the assistance of M. Grange. Z.W., M. Grange, M Gautel and S.R. wrote the manuscript. All authors reviewed the results and commented on the manuscript.

## Materials and Correspondence

Correspondence and requests for materials should be addressed to S.R.

## Methods

### Myofibril isolation

Skeletal myofibrils were prepared from pre-stretched BALB/c mouse psoas muscle fibre bundles as described previously^42^.

Cardiac myofibrils were prepared from left ventricular trabecular strips pre-stretched overnight to a sarcomere length of about 2 µm in rigor buffer (20 mM HEPES pH 7, 140 mM KCl, 2 mM MgCl_2_, 2 mM EGTA, 1 mM DTT, Roche complete protease inhibitor) at 4°C. Left ventricles were cut into ∼1 mm pieces using scalpel blades and homogenised first in rigor buffer with complete protease inhibitors, then resuspended and homogenised 3-4 times in rigor buffer containing 1% (v/v) Triton X-100 essentially as described^45^. Dissociation into myofibril bundles containing 3-5 myofibrils was monitored by microscopy. The concentration of myofibrils was adjusted with complete rigor buffer to ∼5 mg/mL, using an extinction coefficient of myofibrils in 1% (w/v) warm SDS solution of ∼0.7 mL mg^-1^cm^-1^.

### Vitrification of myofibrils and cryo-focused ion beam milling

Myofibrils were frozen on grids by plunge-freezing using a Vitrobot. Generally, 2 µl of myofibril suspension was applied onto the glow-discharged carbon side of Quantifoil R 1.2/1.3 Cu 200 grids. After a 60-second incubation at 13°C, the grids were blotted from the opposite side of the carbon layer for 15 seconds before plunging into liquid ethane. For the dataset aimed at determining the I-band thin filament structure, myofibrils were frozen on Quantifoil R 1/4 Au 200 grids with SiO_2_ film after a longer blotting time of 20 seconds. Frozen grids were clipped into cryo-FIB-specific AutoGrids with marks for grid orientation and a cut-out for low angle FIB-milling.

Clipped grids were transferred into an Aquilos cryo-FIB/SEM dual-beam microscope (Thermo Fisher). Cryo-FIB-milling was performed as previously described^42^. Briefly, the grids were first sputter-coated with platinum and then coated with metalloorganic platinum through a gas-injection-system. The myofibrils were thinned into lamellae in a four-step milling process with an ion beam of decreasing current from 0.5 µA to 50 nA. For the dataset of I-band thin filaments, AutoTEM was used to automatically produce lamellae with thicknesses of 50-200 nm. During auto-milling, an anti-contamination shield replacing the original shutter was inserted to minimise contamination from water deposition^46^.

### Electron cryotomography data acquisition

Grids containing milled lamellae were transferred through a low-humidity glovebox^46^, in order to avoid contamination, into a Titan Krios (Thermo Fisher) transmission electron microscope equipped with a K2 Summit or K3 camera (Gatan) and an energy filter. Projection images were acquired using SerialEM software^47^. Overview images of myofibrils in lamellae were acquired at 6,300 x or 8,400 x nominal magnification to identify locations for high-magnification tilt series acquisition and serve as reference images for batch tomography data acquisition. Tilt series of the skeletal A-band dataset were acquired at 81,000 x nominal magnification (pixel size 1.73 Å, calibrated based on an averaged reconstruction and a crystal structure of myosin (PDB: 3I5G)^48^) with the K2 Summit camera. Tilt series of the cardiac A-band dataset and the skeletal I-band dataset were acquired with the K3 camera, at 42,000 x (pixel size 2.23Å) or 81000 x (pixel size 1.18 Å) nominal magnification, respectively. A dose-symmetric tilting scheme^49^ was used during acquisition with a tilt range of -54° to 54° relative to the lamella plane at 3° increments. A total dose of 130-150 e^-^/Å^2^ was applied to the sample.

### Tomogram reconstruction and automatic filament picking

Individual tilt movies acquired from the microscope were motion-corrected^50^ and combined into stacks for a given tomogram with matched angles using a custom script for subsequent tomogram reconstruction. The combined stacks were then aligned, CTF corrected through strip-based phase flipping and reconstructed using IMOD^51^. In total, three datasets were collected: skeletal (psoas) A-band (171 tomograms), cardiac A-band (24 tomograms) and skeletal I-band (115 tomograms). Tomograms containing wrong field of view, incompletely vitrified ice or lacking visible inherent sarcomere features due to the lamella being too thick (>150 nm) were discarded. Eventually, 48, 24 and 47 tomograms were selected from skeletal A-band, cardiac A-band and skeletal I-band datasets, respectively, for further processing.

#### Automated picking of mouse psoas A-band

The picking of thin filaments within the mouse psoas A-band was performed as previously outlined^42^. Briefly, tomograms were re-oriented so that the thin and thick filaments of the sarcomere lay along the Y axis of the volume and such that XY slices contained the central section of the thin and thick filaments. After applying an equatorial filter as a Fourier mask, thin filaments were recognised and traced from the XZ slices by the TrackMate plugin^52^ in Fiji^53,54^. In total, 183,260 segments of the thin filaments (sub-tomograms) were picked from 48 tomograms with an inter-segment distance of 62 Å. The distance was determined to accommodate two adjacent actin subunits on one strand for further averaging of myosin double-head structure.

#### Automated picking of mouse cardiac A-band

The picking of the mouse cardiac A-band thin filament used a similar approach as described above for skeletal A-band, with the following changes: after the rotation of the tomograms, from the XZ slices, positions of the thin filaments were determined using crYOLO^55^ instead of TrackMate. Manual picking of thin filaments from 28 XZ slices from 6 tomograms were used for initial training in crYOLO. The positions for thin filament in all tomograms were then picked and traced by crYOLO. In total, 202,864 segments of the thin filaments were picked from 24 tomograms with an inter-segment distance of 65 Å.

#### Automated picking of mouse psoas I-band

The picking of the mouse psoas I-band was performed using the latest development version of crYOLO (1.8.0b33) to pick directly on XY slices without any pre-rotation of tomograms. Tomograms were reconstructed using the program Warp after alignment in IMOD, at a down-sampled scale of 8x. Thin filaments were manually picked on 21 XY slices from 4 tomograms. These picked positions were then used to train a model in crYOLO which was used to pick all tomograms. The picked positions were then traced through different XY planes in 3D. In total, 84,937 segments of I-band thin filaments were picked from 47 tomograms with an inter-segment distance of 38 Å.

### Sub-tomogram averaging

#### Skeletal A-band thin filament

Sub-volume averaging of the skeletal A-band thin filaments first followed a previously published approach^42^. Briefly, the determined positions for filaments within the tomograms were used to extract sub-tomograms in RELION^56^ using a box size of 200 voxels (346 Å), which were then projected (central 100 slices) and sorted into good and bad particle classes through 2D classification in ISAC^57^. Good particles were then subject to three-dimensional refinement using a cylindrical reference in RELION, achieving a global resolution (0.143 criterion) of 8.8 Å.

In order to increase resolution, the particles were then subjected to further refinement in the program M^58^. Tilt movies and image stacks were motion-corrected and CTF-estimated within Warp^59^ and new tomograms were reconstructed. The original particle position information obtained from the final step of refinement in RELION was then transformed to match the output geometry of tomograms from Warp. The new particles were extracted in Warp for subsequent averaging in RELION using a 2x down-sampling. After 3D refinement in RELION, the structure of the thin filament was determined to 7.8 Å (Extended Data Fig. 1). The final half-maps and alignment parameters were subjected to M for refinement. The strategy for refinement in M followed previously published regimens^58^. After this refinement, the structure reached a global resolution of 6.7 Å. The core of the thin filament, including actin and nebulin, was masked and reached a resolution of 4.5 Å (Extended Data Figs. 1,2), which was used for model building of actin and nebulin.

#### Skeletal A-band actomyosin and myosin neck domain

In order to resolve the actomyosin structure, including thin filament and a bound myosin double-head, a 3D classification approach similar to the one previously published^42^ was used. After the refinement in M, sub-tomograms were re-extracted and classified in RELION. Classes were translated and rotated to a common double-head configuration and re-refined in M. The final reconstruction at a resolution of 6.6 Å was used for model building of myosin heavy chain. In order to increase the resolution of the myosin neck domains (predominantly the ELC and RLC), the sub-tomograms were first re-centred towards ELC and re-refined in RELION with a mask containing only myosin. This resulted in an ELC-centred myosin double-head structure with a resolution of 8.9 Å (Extended Data Fig. 1). Afterwards, the sub-tomograms were further re-centred towards RLC and re-refined with a smaller mask containing ELCs and RLCs to reconstruct a structure of RLC-centred myosin double-head with a resolution of 9 Å. The ELC- and RLC-centred myosin double-head maps were used for rigid-body docking of ELC and RLC models, respectively.

#### Cardiac A-band thin filament, actomyosin and myosin neck domain

Sub-volume averaging of the cardiac A-band thin filament followed the same strategy as for the A-band of skeletal muscle and resulted in a structure of thin filament resolved to a global resolution of 8 Å, with the core of thin filament (actin) resolved to 6.3 Å (Extended Data Fig. 3a,b). Structures of cardiac actomyosin and myosin neck domain were determined using the same classification and re-centring approach as for skeletal structures, resulting in resolutions of 7.7 Å and 12 Å, respectively.

#### Skeletal I-band thin filament

Sub-volume averaging of the skeletal I-band was performed largely as described for the A-band structures, except for using helical symmetry (twist -167.4°, rise 28.8 Å) during the initial refinement in RELION to reduce alignment error due to the missing wedge artefacts. The final structure of the I-band thin filament was determined to a global resolution of 9.4Å, with the core (actin and nebulin) resolved to 7.4 Å (Extended Data Fig. 3c,d).

### Model building of actin, nebulin and myosin heavy chain

To reduce the risk of over-refinement and account for the heterogenous resolution of our structures, several density maps, which were masked to different areas, filtered to nominal or local resolution as determined by SPHIRE^60^ and sharpened using various B-factors, were used for model building. In addition, density modified maps were calculated from half maps providing the reported nominal resolution^61^.

An initial model for actin was generated by homology modelling using Modeller^62^ in Chimera based on a previous atomic model (PDB: 5JLH^63^, chain A) and a sequence alignment from Clustal Ω. The unresolved N-terminus of actin (aa 1-6) was removed and Mg^2+-^ADP was added from PDB 5LJH. HIS 73 was replaced by HIC (4-methyl-histidine) and regularized in Coot^64^. A pentameric composite model was assembled by rigid-body fitting in Chimera including an initial model of nebulin (see below). Model building was performed in ISOLDE^65^ in ChimeraX^66^. A total of four density maps were loaded (filtered to nominal resolution and sharpened with B-factors -70 and -150; filtered to local resolution and sharpened to B-factor - 100; and the density modified map). Only the central actin chain and residues in close contact were included in the simulation and rebuilt. Unresolved side chains are in the most likely positions. After a first pass through the complete molecule, Ramachandran and rotamer issues were addressed locally. Based on the refined central chain, the composite actin-nebulin pentamer was updated. Hydrogens were removed and the resulting model was real-space refined against the map filtered to nominal resolution in Phenix^67^. To avoid large deviations from the input model, the ISOLDE model was used as a reference, while local grid search, rotamer and Ramachandran restraints were deactivated. The actin model was further improved by a second round of model building in ISOLDE.

Modeling of nebulin was performed in analogy. An initial poly-alanine model for nebulin was built manually in Coot based on the density of the central repeat (4.5 Å resolution). In order to cover the connection between two nebulin repeats, a peptide of 56 aa (instead of 35 aa) was initially built. The density corresponding to residue 22 was consistent with a consensus tyrosine residue and thus tyrosine was used instead of alanine. A segmented post-processed map (filtered to nominal resolution and sharpened with B-factor -70) was loaded for further modelling in ISOLDE. Secondary structure and rotamer restraints were applied where appropriate. Based on the resulting model, a continuous model of nebulin was created by first cutting the model to 35 aa and rigid-body fitting into the density. The termini of three consecutive nebulin chains were then manually connected. To address geometry issues due to the connection, the combined nebulin chain was real-space refined in Phenix against the segmented map and subjected to another round of refinement in ISOLDE.

Refined models of actin and nebulin were finally combined into one pentameric model. Minor adjustments to the orientation of side chains were done in Coot where necessary. The composite model was real-space refined against the 4.5 Å-resolution map filtered to local resolution (B-factor -100) in Phenix using the same settings as before.

An initial model of the actin-nebulin-tropomyosin-myosin (actomyosin) complex was assembled from the refined actin-nebulin model, a homology model of myosin^48^ (PDB: 3I5G, chain A), and a polyalanine model of tropomyosin^63^ (PDB: 5JLH chains J and K) using rigid-body fitting. Only the heavy chain of myosin (up to residue 788) was modelled. After addition of hydrogens, the central myosin chain was refined in ISOLDE using four segmented density maps of actomyosin, as described for actin. All applicable secondary structure restraints and many rotamer restraints were applied. Manual building was started from the acto-myosin interface, as it is best resolved. Unresolved residues including loop I (aa 207-215), loop 2 (626-643) and the N-terminus (1-11) were removed. After deletion of hydrogens, the resulting atomic model was real-space refined in Phenix and further improved by a second round of refinement in ISOLDE. The refined atomic model of the central myosin chain, was finally used to assemble an updated composite model of the actomyosin complex. This model was addressed to a final round of real-space refinement in Phenix against a 6.6 Å-resolution density map filtered to nominal resolution (B-factor -75) using the same settings as before, but with both Ramachandran and Rotamer restraints applied.

As the resolution was not sufficient to reliably model Mg^2+^ ions, they were replaced with the ones from PDB 5JLH by superposition of actin subunits. The final atomic models of actin-nebulin and actomyosin complexes were assessed by Molprobity^68^ and EMRinger^69^ statistics (Extended Data Table 1).

### Rigid-body docking of myosin light chains

As the density for the C-terminus of the myosin heavy chain lever arm as well as the ELC and RLC is of insufficient quality for reliable model building with refinement, rigid-body docking of previously published structural models^48^ (PDB: 3I5G) was performed. First, the ELC model together with the ELC-binding lever arm helix (aa 785-802 in PDB 3I5G) were docked into both myosin ELC densities in the ELC-centred myosin double-head map (8.9 Å, B-factor -500) in Chimera. Then, the RLC model together with RLC-binding HC helices (aa 809-839) were docked into the RLC density of the lower myosin head in the RLC-centred myosin double head map (9 Å, B-factor -300). For the RLC of the upper head, the C-lobe of RLC (together with HC helix aa 809-824) and the N-lobe of RLC (together with HC helix aa 826-839) were docked separately into a segmented map of upper myosin RLC (Fig. S9f). The maps of actomyosin, ELC-centred myosin double-head and RLC-centred myosin double-head were aligned in Chimera in order to unify the coordinate system of all models. In the end, a final homology model was calculated based on these initial models and the sequences of mouse myosin heavy chain and light chains from mouse fast muscle using SWISS-MODEL^70^. In order to compare the difference of RLC-RLC interface between active and inactive myosin, this model was compared to previous structures of myosin IHM^43^ (PDB: 6XE9) through aligning the lower head RLC from our model to the free head RLC from IHM in Chimera.

### Sequence analysis of nebulin and troponin T

As a defined boundary on nebulin sequence between A-band and I-band is not present, the nebulin sequence of M1-8 and the entire super repeat region (Extended Data Fig. 2a) from mouse (Uniprot: E9Q1W3) was considered as the A-band nebulin sequence and divided into 176 simple repeats (M1-162) through placing the SDxxYK motif at position 18-23. Multiple sequence alignment was performed using ClustalW^71^ with gaps disabled (Extended Data Fig. 6a) and visualised in WebLogo^72^. Secondary structure of each simple repeat was predicted using RaptorX-Property^73^. Probability values for being α-helix at each residue position were averaged and used for Fig. 2 and Extended Data Fig. 6. To estimate relationship between the charge of the amino acid at position 15 and 21 a Bayesian multi-nominal regression was performed. For both positions and for each of the 176 sequences, the amino acid type was assigned a number, representing one of four categories which are 1 (Positive), 2 (Negative), 3

(Hydrophobic) and 4 (Other). With this, the categorical variables 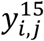and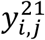were constructed, representing the category *i* at position 15 and 21 for sequence *j* respectively. The hierarchical Bayesian model was then modelled the following way and fitted with Stan^74^:

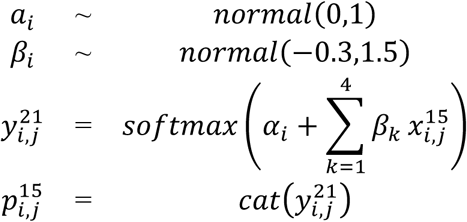

Where *a*_*i*_ is the intercept, *β*_*i*_ the regression coefficients and 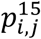 the probability of seeing category *i* in sequence *j*. The variable 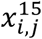 is an indicator variable which is one when sequence *j* at position 15 is of class *i*. The priors for *a*_*i*_ and *β*_*i*_ were chosen in a way to that the prior predictive distribution of 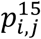 has mean probability for each class of 0.18. After fitting, 1000 samples were drawn from the posterior distribution for each possible state of category of amino acid 21 (Extended Data Fig. 7b).

Troponin T linker sequence was from mouse fast skeletal muscle troponin T (UniProt: Q9QZ47-1) after sequence alignment with the sequence of the missing segment of troponin T (R151-S198) in PDB: 6KN8^35^. Hydrophobicity score (Extended Data Fig. 8g) was calculated through ProtScale^75^ using the scale from Abraham & Leo^76^.

## Data availability

Cryo-ET structures and representative tomograms are deposited in the Electron Microscopy Data Bank under accession numbers EMD-XXXXX, EMD-XXXXX, EMD-XXXXX, EMD-XXXXX, EMD-XXXXX, EMD-XXXXX, EMD-XXXXX, EMD-XXXXX, EMD-XXXXX, EMD-XXXXX, EMD-XXXXX. The atomic models are deposited in Protein Data Bank under accession numbers XXXX, XXXX and XXXX. Other structures and EM density maps used in this study are available in PDB: 3I5G, 6KN8, 6XE9 and in EMDB: EMD-0729. The datasets generated and analysed during the current study are available from the corresponding author on reasonable request.

---

**Extended Data Fig. 1.**
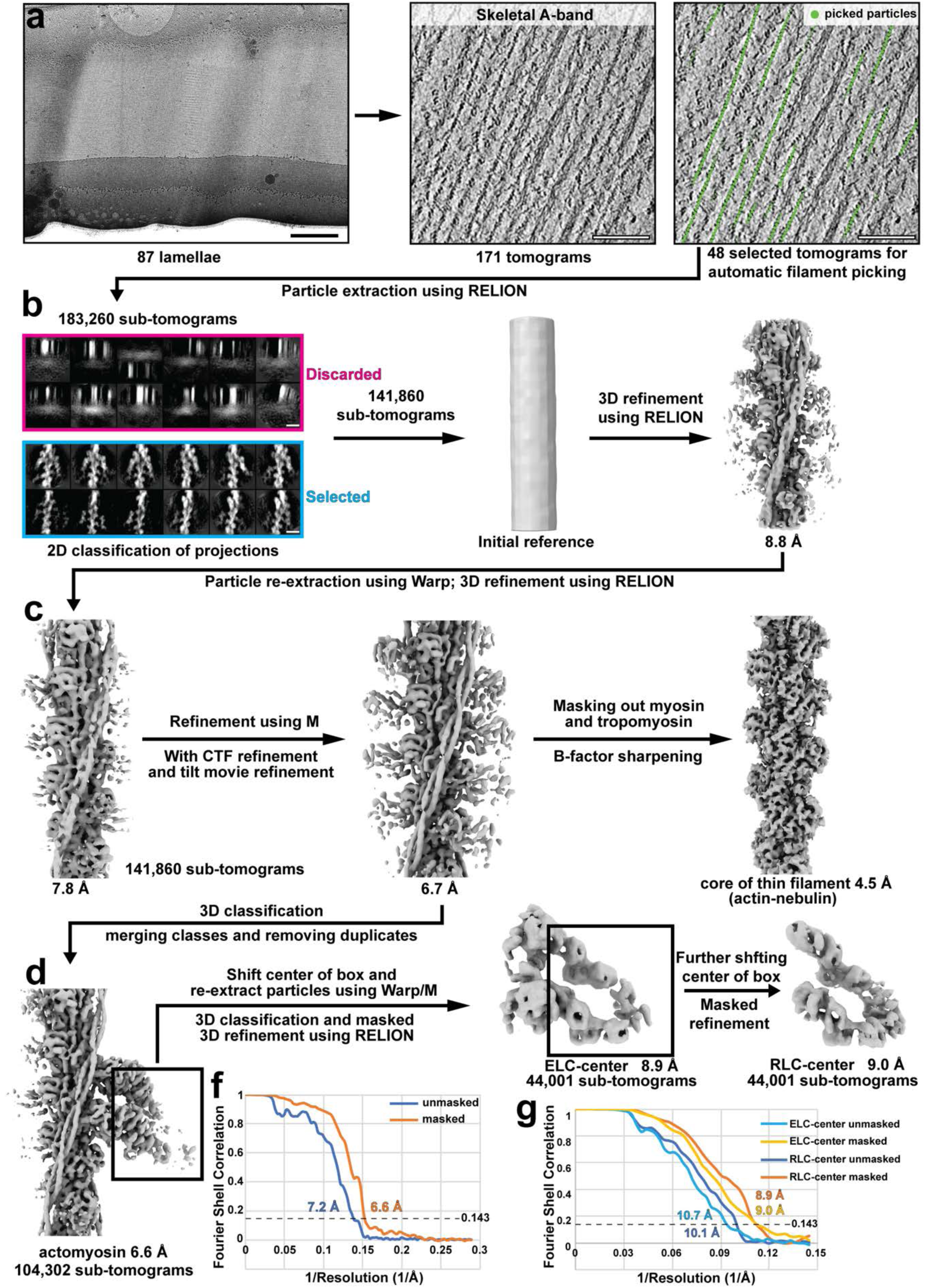
Cryo-FIB-ET and processing workflow of A-band thin filament structures from mouse psoas muscle. **a**. Examples of a lamella of mouse psoas myofibrils and a slice of tomogram depicting the sarcomeric A-band, where the thin filaments are picked automatically. Scale bars: 1 µm (left), 100 nm (right). **b**. Cleaning of particles using 2D classification of projection images and initial refinement using RELION. Scale bar: 10 nm. **c**. Improving the resolution of the thin filament structure using the Warp-M packages. **d**. Actomyosin structure (including a thin filament and two myosin heads) and structures of the myosin neck domain obtained through further classification and re-centring of particle boxes. **f**. Gold-standard FSC curve of the actomyosin complex. **g**. Gold-standard FSC curve of the ELC and RLC centred myosin double head structures.

**Extended Data Fig. 2.**
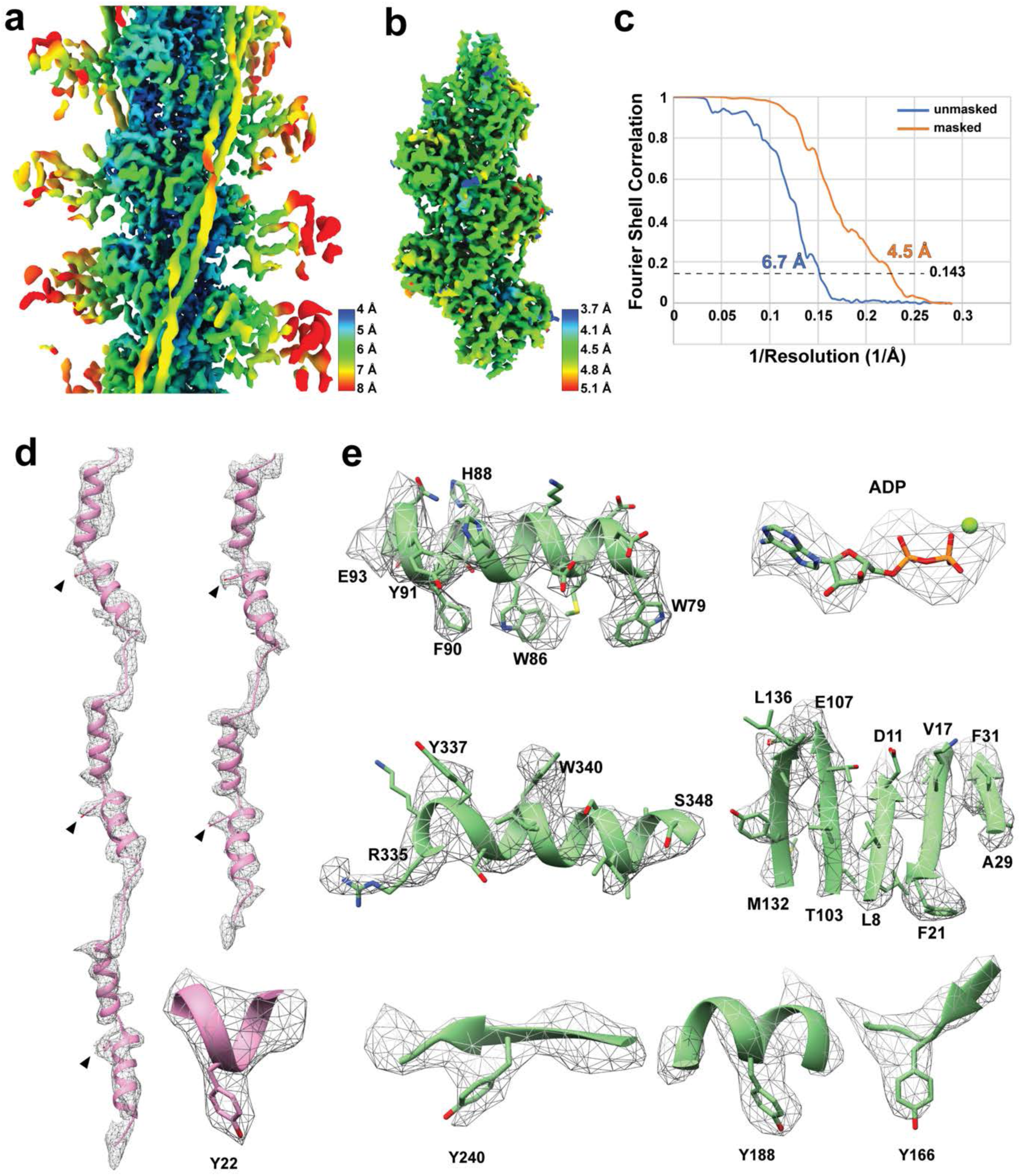
EM density map and structural model of actin and nebulin obtained from sub-tomogram averaging. **a**. Local resolution estimation of the unmasked thin filament. The map is filtered to local resolution. **b**. Local resolution estimation of the masked core of the thin filament including actin and nebulin. **c**. Gold-standard FSC curve of the actin-nebulin structure. The unmasked and masked structures correspond to the maps in (**a**) and (**b**), respectively. **d**. Nebulin model and corresponding cryo-EM density map. The conserved tyrosine residues are marked with arrow heads. A zoom-in view of an example of tyrosine 22 and corresponding density is shown. **e**. Examples of side chain density visible in the actin portion of the cryo-EM density map. A few tyrosine residues are selected for comparison with the tyrosine side chain densities in nebulin.

**Extended Data Fig. 3.**
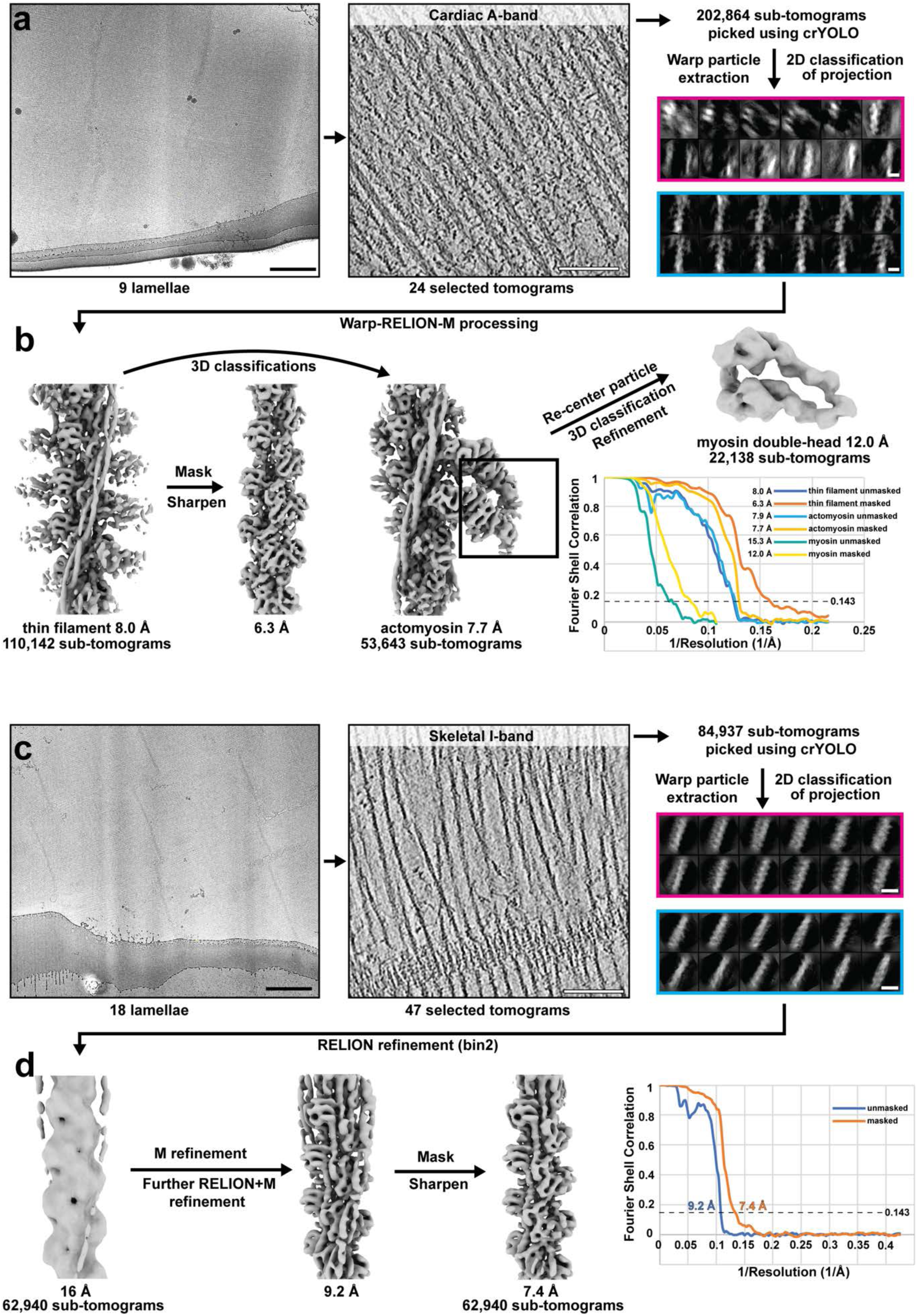
Cryo-FIB-ET and processing workflow of A-band thin filament structures from mouse cardiac muscle and I-band thin filament structures from mouse psoas muscle. **a**. Examples of a lamellae of mouse cardiac myofibrils, a slice through a tomogram of mouse cardiac sarcomere A-band and particle-cleaning using 2D classification. **b**. EM density maps and gold-standard FSC curves of the thin filament, actomyosin and re-centred myosin double-head structures. The processing workflow is similar to the processing of mouse skeletal A-band structures. **c**. Examples of a lamellae of mouse psoas myofibrils, a slice through a tomogram of mouse skeletal sarcomere I-band (also depicting a Z-disc at the bottom) and particle-cleaning using 2D classification. **d**. Processing workflow of the I-band thin filament and gold-standard FSC curve of the structure. **e**. Scale bar: 1 µm (lamella), 100 nm (tomogram), 10 nm (2D classes)

**Extended Data Fig. 4.**
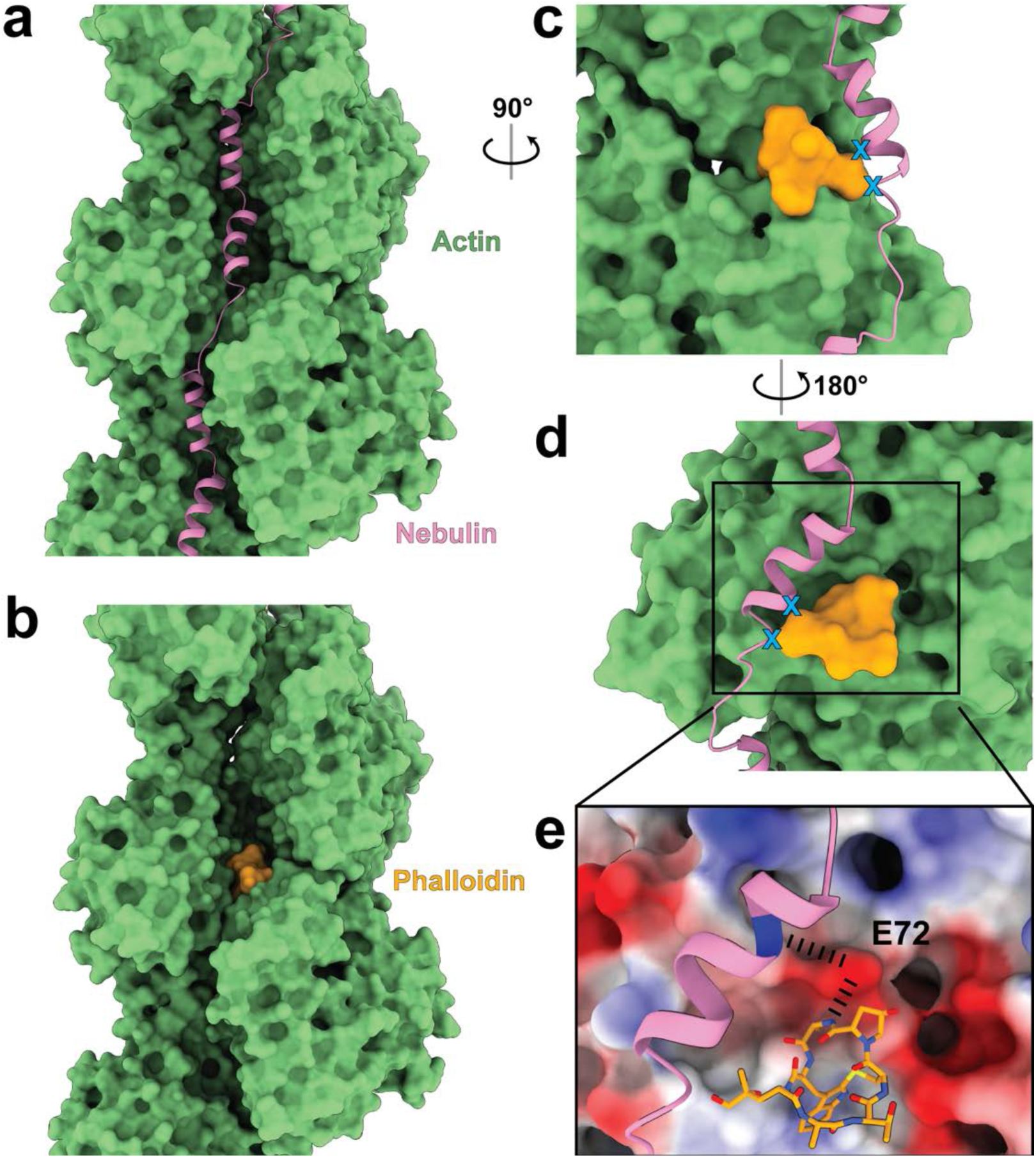
Comparison of nebulin and phalloidin binding sites on the actin filament. **a-b**. Position of nebulin and phalloidin (from PDB: 6T1Y) on the actin filament, respectively. **c**,**d**. Comparison of the position of both nebulin and phalloidin from different views. Steric clashes would happen at the end of H2 of nebulin if both were present at the same time (marked by blue crosses). **e**. E72 on actin is involved in both forming a hydrogen bond with phalloidin and electrostatic interactions with nebulin.

**Extended Data Fig. 5.**
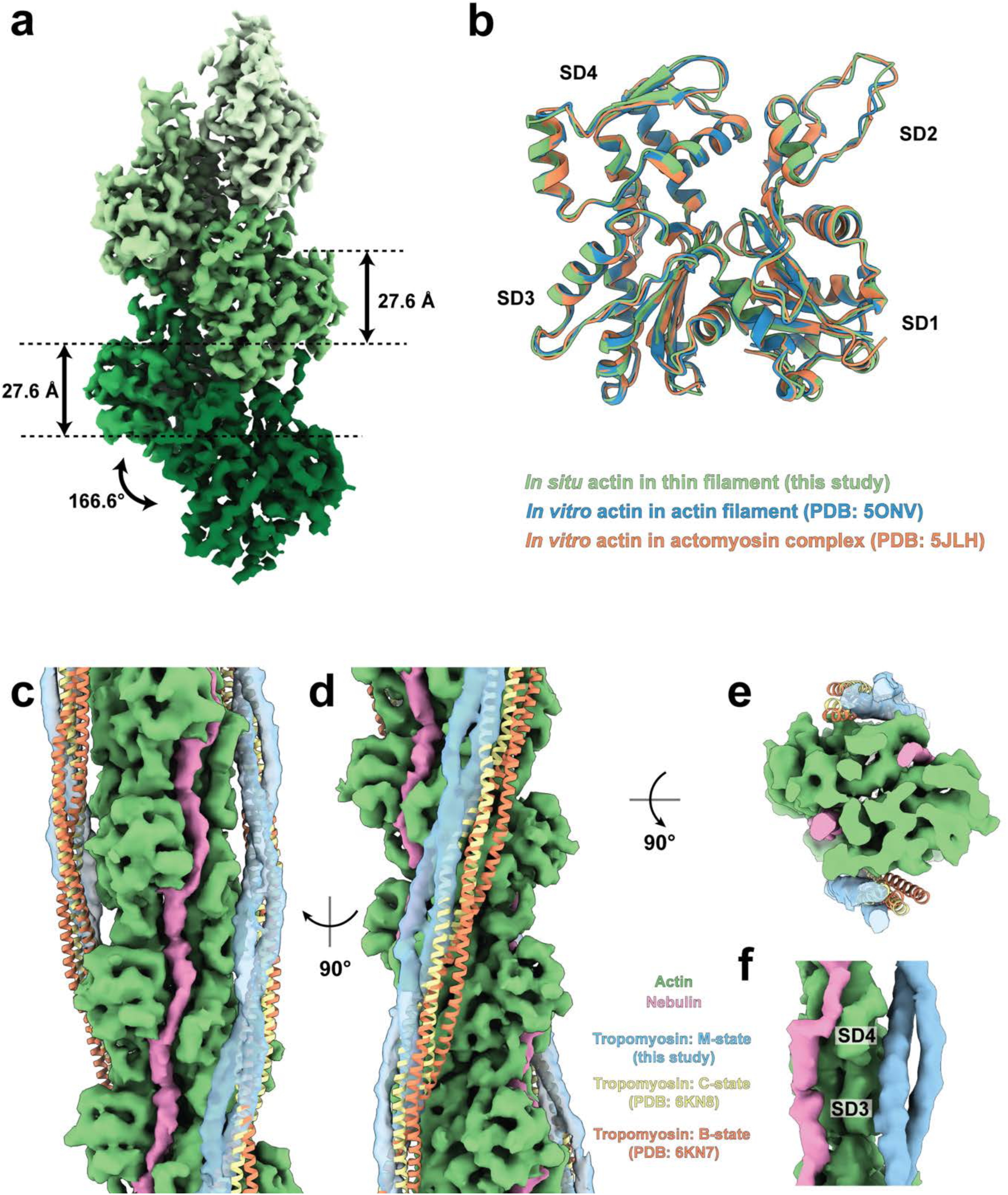
Actin in a thin filament and different tropomyosin states on a thin filament. **a**. Helical parameters of F-actin determined within a thin filament. **b**. Comparison of the structure of actin from different filamentous structures. **c-e**. Different views depicting a thin filament including nebulin and different states of tropomyosin. **f**. Zoom-in view of nebulin, tropomyosin and actin depicting the physical separation between nebulin and tropomyosin by the SD3 and 4 of actin subunits.

**Extended Data Fig. 6.**
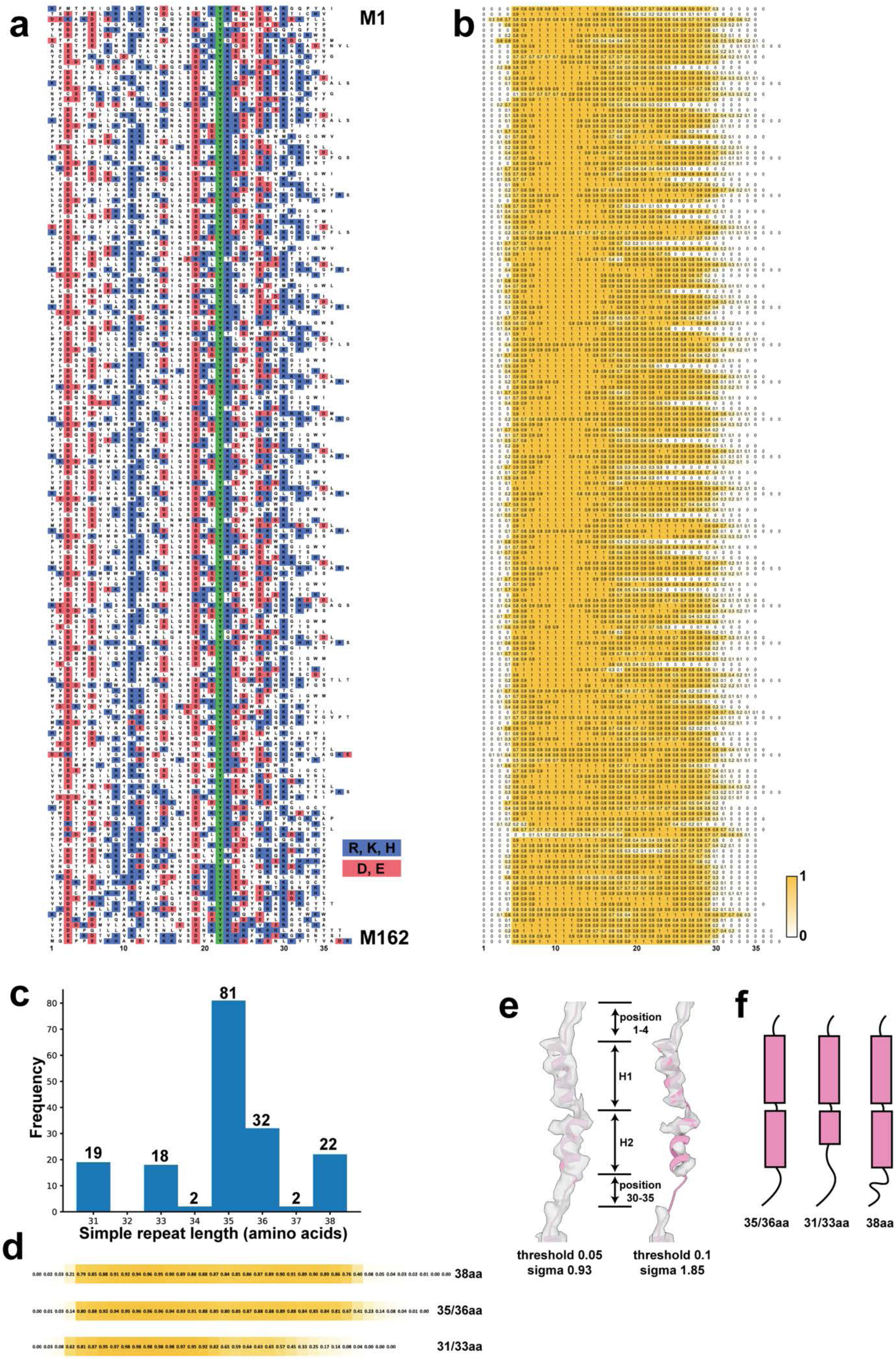
Sequence, secondary structure prediction and length of individual nebulin simple repeats. **a**. Sequence alignment of nebulin repeats M1-M162. Positively and negatively charged residues are highlighted in blue and red, respectively. The fully conserved tyrosine at position 22 is highlighted in green. **b**. Secondary structure prediction of repeats M1-M162, coloured based on the probability of an α-helix. **c**. Histogram of the length of simple repeats. **d**. Averaged secondary structure prediction of normal (35/36 aa), short (31/33 aa) and long (38 aa) simple repeats. **e**. Cryo-EM density map of one nebulin repeat at low and high surface threshold. **f**. Schematic model of the nebulin simple repeats with different numbers of amino acids but the same physical length. α-helices are represented in magenta rectangles. Loops are shown as lines.

**Extended Data Fig. 7.**
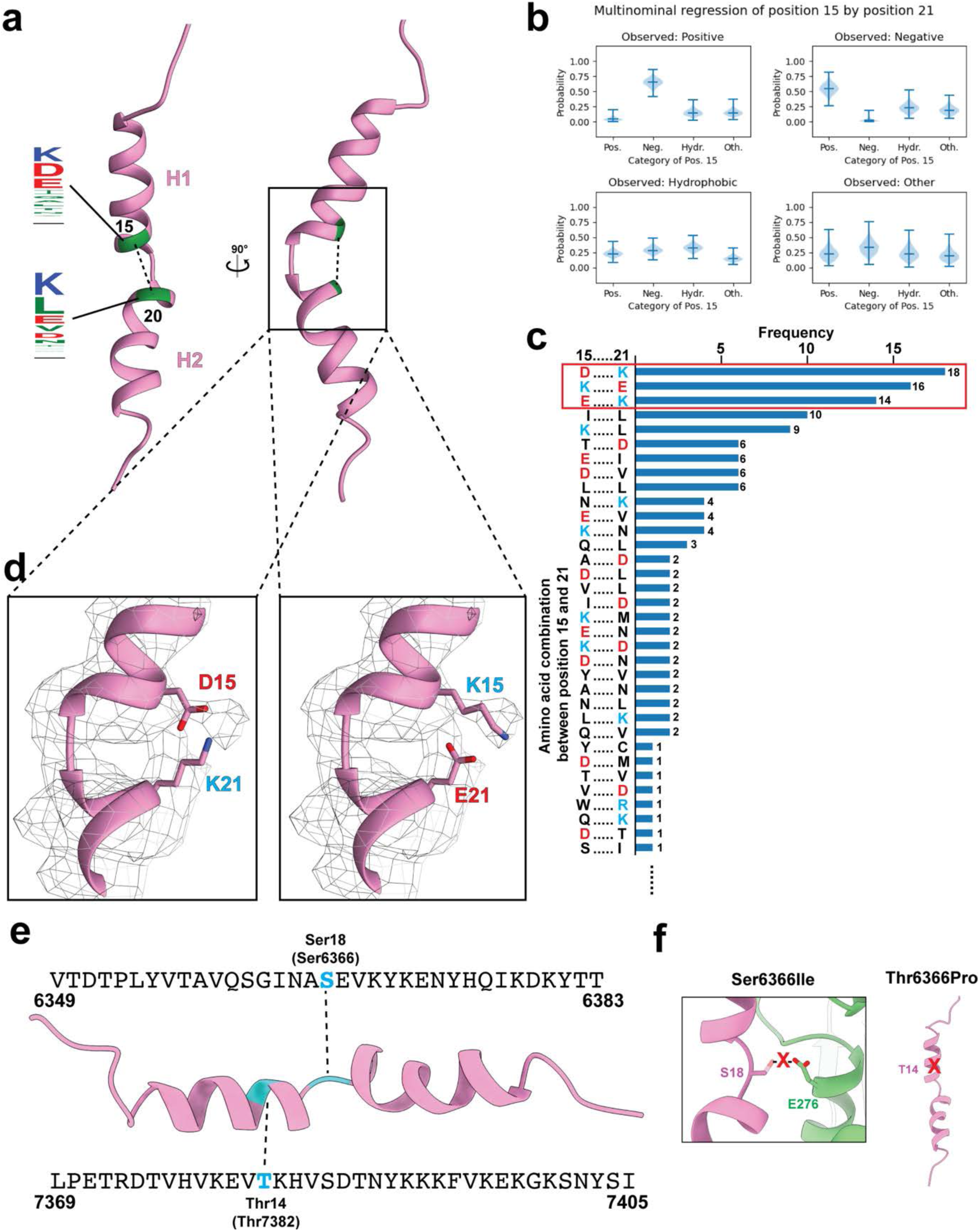
Intra-nebulin interactions between position 15 and 21 and location of pathogenic missense mutations on a simple repeat. **a**. Location of residue number 15 and 21 on a simple repeat including the possible amino acids at these two positions. Larger amino acid letter represents higher occurrence. **b**. Violin plot of Bayesian multinominal regression depicting the conditional probability of a certain amino acid type on position 15 given the amino acid residue type on position 21. The median, minimum and maximum estimated probability values for each category are shown. The shaded area shows the probability density of the model within the respective category. **c**. Histogram of amino acid combinations of position 15 and 21. Charged amino acids are coloured in blue (positive) and red (negative). Other combinations with only one occurrence are not shown. **d**. Weak side chain densities at position 15 and 21. Fitted structural models are mutated to the two most populated amino acid combinations at position 15 and 21 showing possible side chain conformations. **e**. Location of the two founder mutations of nemaline myopathies in the Finnish population, S6366I and T7382P, on a simple repeat. **f**. Potential disruption of a hydrogen bond between nebulin and E276 on actin due to the mutation S6366I and potential disruption of the H1 helix structure due to the mutation T7382P.

**Extended Data Fig. 8.**
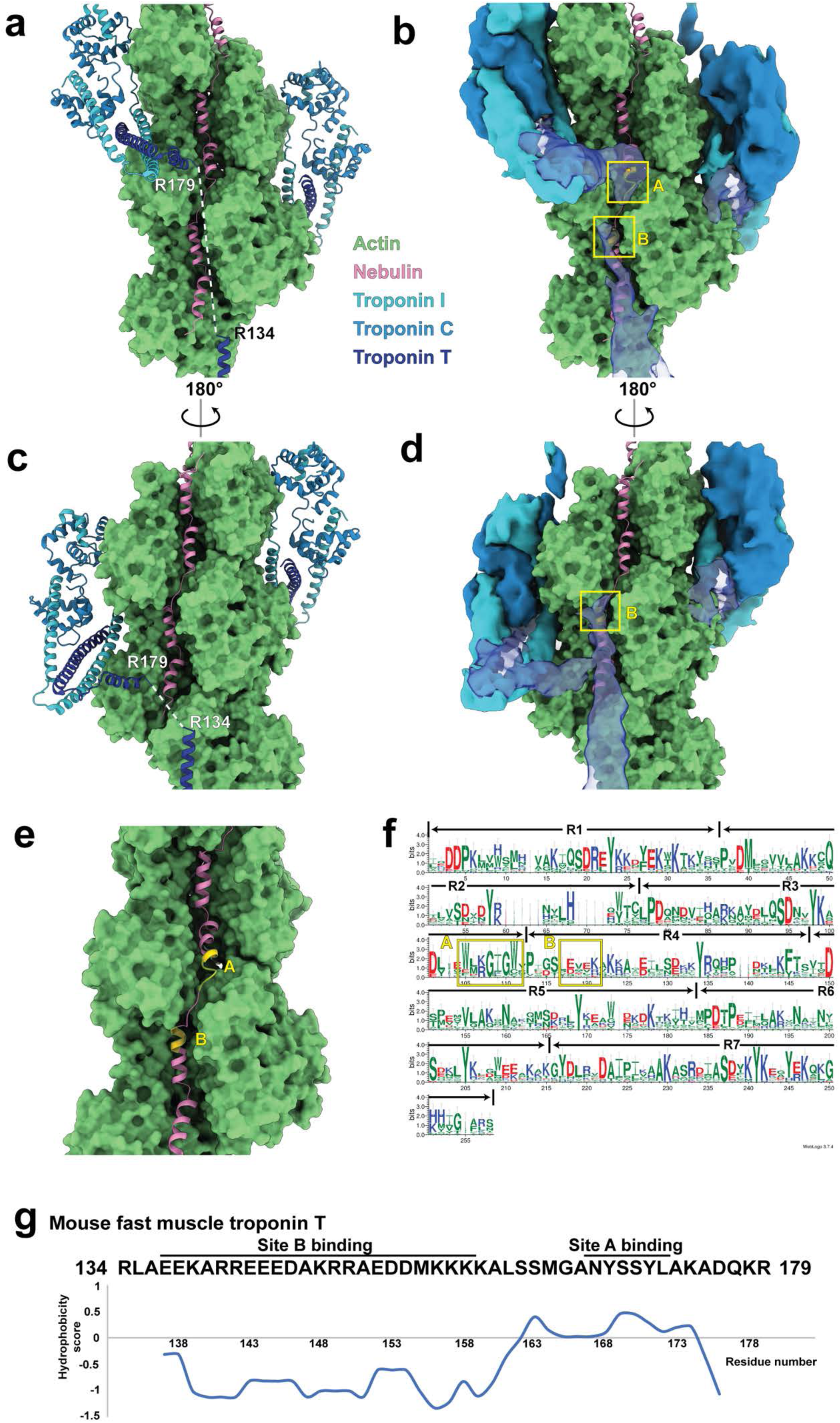
Interactions between nebulin and troponin. **a-d**. The actin-nebulin complex superimposed with the structural models (**a**,**c**) and cryo-EM densities (**b**,**d**) of the troponin complex (PDB:6KN8; EMD-0729). The linker region between R134 and R179 (corresponding to R151 and S198 in the original model) is missing in the structural model. Two contact sites between nebulin and troponin T are marked in yellow as A and B. **e**. The two troponin T binding sites are highlighted in yellow on nebulin. **f**. Graphical representation of sequence alignment of all nebulin super repeats (each super repeat contains the simple repeats R1-R7). A larger amino acid symbol corresponds to a greater occurrence at a certain position. The troponin binding sites A and B are marked corresponding to the WLKGIGW and ExxK motif. **g**. Hydrophobicity of the linker in mouse fast muscle troponin T (TNNT3). Potential regions that can bind site B and site A are marked.

**Extended Data Fig. 9.**
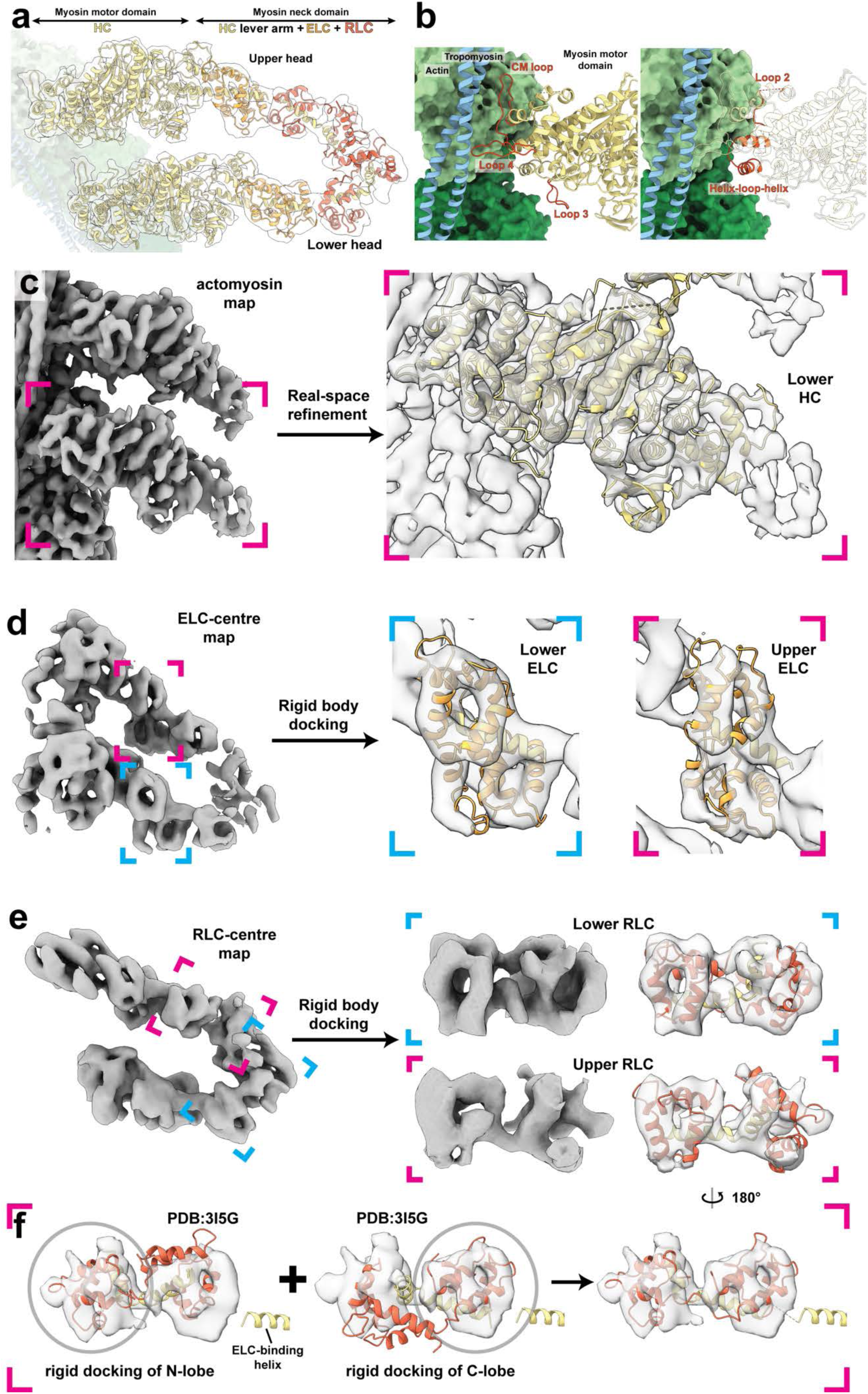
Model building of myosin heads. **a**. Composite EM density map and a complete structural model of a myosin double-head when bound to a thin filament. **b**. Actin-myosin interaction interface. Loops on myosin involved in the interaction with actin are highlighted in red. **c**. Cryo-EM density map of myosin motor domain with structural models (yellow) refined into the map. **d**. Cryo-EM density map of myosin heads with a focus on the ELC. Myosin ELC (orange) and HC LH2 (yellow) from PDB 3I5G were docked into the map as rigid bodies. **e**. EM density map of myosin heads with a focus on the RLC. The lower RLC density was directly fitted with the myosin RLC model (red; with HC LH3-4, yellow) from PDB 3I5G. The upper RLC density was fitted with a modified model of RLC and HC LH3-4, as shown in (**d**). **f**. Separate rigid body docking of the N-lobe (red; with HC LH4, yellow) and the C-lobe (red; with HC LH3, yellow) of the RLC in the upper head. The final structural model was obtained combining the two docked models.

**Extended Data Fig. 10.**
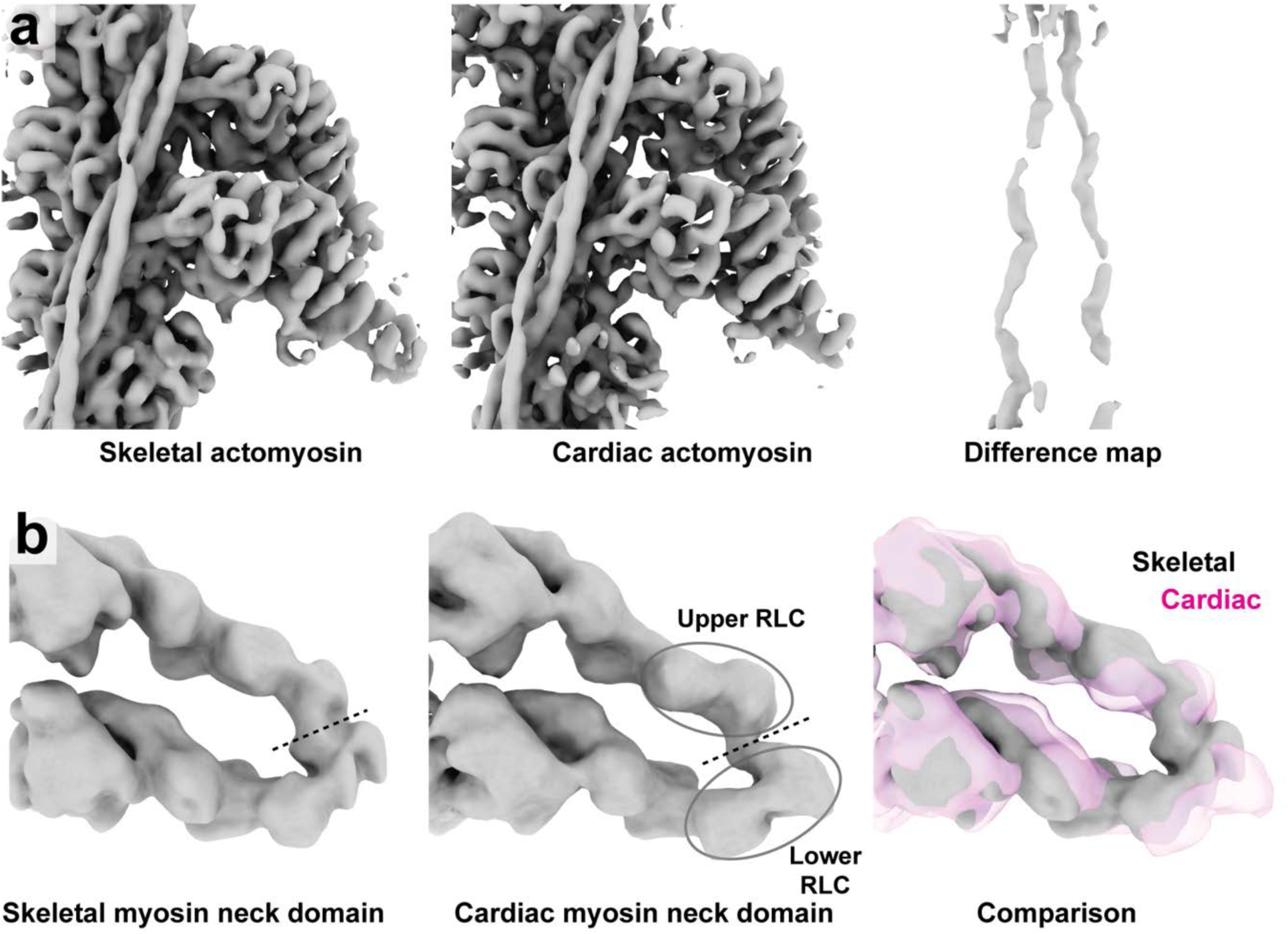
Comparison between cardiac and skeletal actomyosin structures. **a**. EM density maps of skeletal and cardiac actomyosin, both filtered to 7.8 Å, as well as the difference maps between them. The densities in the difference map correspond to nebulin. **b**. EM density maps of skeletal and cardiac myosin double head, with a focus on the neck domain, both filtered to 15 Å. Dotted lines mark the interfaces between upper and lower RLCs. The comparison between the skeletal and cardiac map is shown on the right.

**Extended Data Fig. 11.**
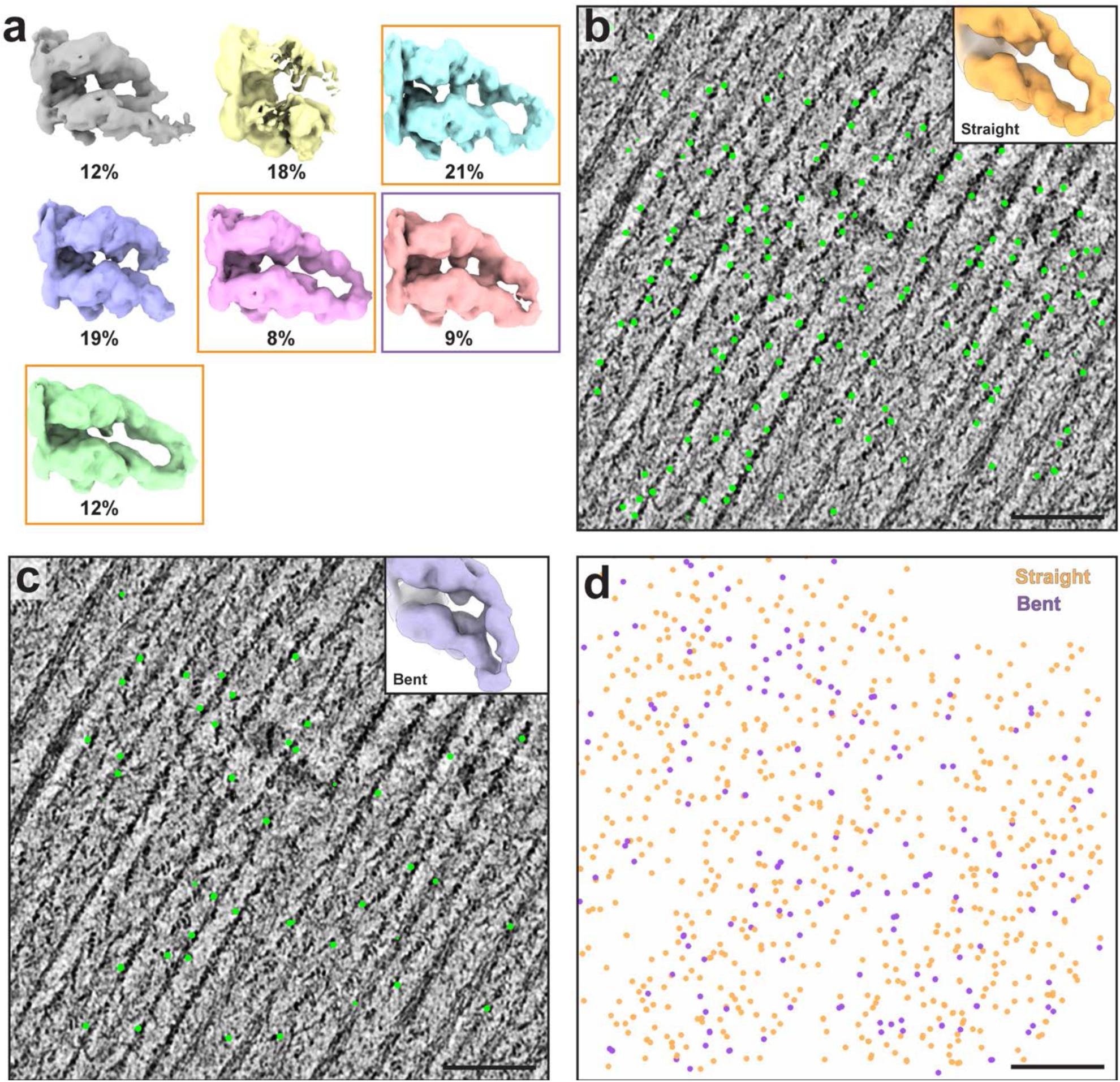
Distribution of the two conformations of myosin double-head. **a**. 3D classification of a myosin double-head. Classes marked in orange and purple are selected as the straight and bent form for further refinement, respectively. **b**. Distribution of myosin double-head in the straight conformation in a tomogram. A slice image of the tomogram with a thickness of 14 nm and particles within it (orange) are shown. **c**. Distribution of myosin double-head in the bent conformation in the same slice of tomogram as (**b**). Particle positions are represented in purple dots. **d**. Distribution of both straight and bent forms in the entire tomographic volume. Scale bars: 100 nm.

**Extended Data Table 1.** Data collection, refinement and model building statistics.

|  | Skeletal A-band |  |  |  | Cardiac A-band |  |  | Skeletal I-band |
| --- | --- | --- | --- | --- | --- | --- | --- | --- |
| Microscopy |  |  |  |  |  |  |  |  |
| Microscope | Titan Krios |  |  |  | Titan Krios |  |  | Titan Krios |
| Voltage (kV) | 300 |  |  |  | 300 |  |  | 300 |
| Camera | Gatan K2 Summit |  |  |  | Gatan K3 |  |  | Gatan K3 |
| Slit width (eV) | 20 |  |  |  | 20 |  |  | 20 |
| Pixel size (Å) | 1.73 |  |  |  | 2.32 |  |  | 1.18 |
| Defocus range (µm) | 2.4-5.0 |  |  |  | 3.2-5.0 |  |  | 2.5-5.5 |
| Tilt range<sup>a</sup> | -54°/+54° (3°) |  |  |  | -54°/+54° (3°) |  |  | -54°/+54° (3°) |
| Tilt scheme | Dose-symmetric |  |  |  | Dose-symmetric |  |  | Dose-symmetric |
| Total dose (e<sup>-</sup>/Å<sup>2</sup>) | 130-150 |  |  |  | 130-150 |  |  | 130-150 |
| Number of lamellae | 87 |  |  |  | 9 |  |  | 18 |
| Number of tomograms<sup>b</sup> | 48 (171) |  |  |  | 24 (24) |  |  | 47 (115) |
| 3D refinement statistics | Actin-nebulin | Acto-myosin | Myosin (ELC-centred) | Myosin (RLC-centred) | Actin | Acto-myosin | Myosin (neck-domain) | Actin-nebulin |
| Number of sub-tomograms | 141,860 | 104,302 | 44,001 | 44,001 | 110,142 | 53,643 | 22,138 | 62,940 |
| Symmetry | C1 | C1 | C1 | C1 | C1 | C1 | C1 | C1, Helical<sup>d</sup><br>(-167.4°, 28.8Å) |
| Resolution<sup>c</sup> (Å) | 4.5<br>(6.7) | 6.6<br>(7.2) | 8.9<br>(10.7) | 9.0<br>(10.1) | 6.3<br>(8.0) | 7.7<br>(7.9) | 12.0<br>(15.3) | 7.4<br>(9.2) |
| B-factor | -100 | -75 | -500 | -300 | -300 | -200 | -300 | -250 |
| EMDB deposition | EMD-XXXX | EMD-XXXX | EMD-XXXX | EMD-XXXX | EMD-XXXX | EMD-XXXX | EMD-XXXX | EMD-XXXX |
| Atomic model statistics |  |  |  |  |  |  |  |  |
| Non-hydrogen atoms | 15,550 | 29,589 | Homology model |  |  |  |  |  |
| Cross correlation masked | 0.72 | 0.82 |  |  |  |  |  |  |
| Molprobity score | 1.75 | 1.97 |  |  |  |  |  |  |
| Clashscore | 17.88 | 30.61 |  |  |  |  |  |  |
| EMRinger score | 0.86 | 0.20 |  |  |  |  |  |  |
| Bond RMSD (Å) | 0.010 | 0.007 |  |  |  |  |  |  |
| Angle RMSD (°) | 1.535 | 1.415 |  |  |  |  |  |  |
| Rotamer outliers (%) | 0.00 | 0.00 |  |  |  |  |  |  |
| Ramachandran favored (%) | 100.00 | 99.69 |  |  |  |  |  |  |
| Ramachandran outliers (%) | 0.00 | 0.20 |  |  |  |  |  |  |
| CaBLAM outliers (%) | 2.15 | 2.33 |  |  |  |  |  |  |
| PDB deposition | PDB: XXXX | PDB: XXXX | PDB: XXXX |  |  |  |  |  |

<sup>a</sup>Tilt angle is relative to the pre-tilt of each lamella. In parenthesis is the tilt angle increment.
<sup>b</sup>In parenthesis is the initial number of tomograms
<sup>c</sup>Resolution is based on 0.143 FSC threshold. In parenthesis is the global resolution without mask applied.
<sup>d</sup>Helical refinement was used for initial refinement in RELION. No helical symmetry was applied during local refinement in RELION and M.

## References

1. Yuen, M. & Ottenheijm, C. A. C. Nebulin: big protein with big responsibilities. J. Muscle Res. Cell Motil. 41, 103–124 (2020).

2. Pappas, C. T., Krieg, P. A. & Gregorio, C. C. Nebulin regulates actin filament lengths by a stabilization mechanism. J. Cell Biol. 189, 859–870 (2010).

3. Witt, C. C. et al. Nebulin regulates thin filament length, contractility, and Z-disk structure in vivo. EMBO J. 25, 3843–3855 (2006).

4. Tonino, P. et al. The giant protein titin regulates the length of the striated muscle thick filament. Nat. Commun. 8, 1–10 (2017).

5. Wang, K. & Williamson, C. L. Identification of an N2 line protein of striated muscle. Proc. Natl. Acad. Sci. U. S. A. 77, 3254–3258 (1980).

6. Wang, K. & Wright, J. Architecture of the sarcomere matrix of skeletal muscle: Immunoelectron microscopic evidence that suggests a set of parallel inextensible nebulin filaments anchored at the Z line. J. Cell Biol. 107, 2199–2212 (1988).

7. Romero, N. B., Sandaradura, S. A. & Clarke, N. F. Recent advances in nemaline myopathy. Curr. Opin. Neurol. 26, 519–526 (2013).

8. Sewry, C. A., Laitila, J. M. & Wallgren-Pettersson, C. Nemaline myopathies: a current view. J. Muscle Res. Cell Motil. 40, 111–126 (2019).

9. Ryan, M. M. et al. Nemaline myopathy: A clinical study of 143 cases. Ann. Neurol. 50, 312–320 (2001).

10. Moncman, C. L. & Wang, K. Nebulette: A 107 kD nebulin like protein in cardiac muscle. Cell Motil. Cytoskeleton 32, 205–225 (1995).

11. Burgoyne, T., Muhamad, F. & Luther, P. K. Visualization of cardiac muscle thin filaments and measurement of their lengths by electron tomography. Cardiovasc. Res. 77, 707–712 (2008).

12. Kolb, J. et al. Thin filament length in the cardiac sarcomere varies with sarcomere length but is independent of titin and nebulin. J. Mol. Cell. Cardiol. 97, 286–294 (2016).

13. Labeit, S. & Kolmerer, B. The Complete Primary Structure of Human Nebulin and its Correlation to Muscle Structure. Mol. Biol. 308–315 (1995).

14. Donner, K., Sandbacka, M., Lehtokari, V. L., Wallgren-Pettersson, C. & Pelin, K. Complete genomic structure of the human nebulin gene and identification of alternatively spliced transcripts. Eur. J. Hum. Genet. 12, 744–751 (2004).

15. McElhinny, A. S., Kolmerer, B., Fowler, V. M., Labeit, S. & Gregorio, C. C. The N-terminal end of nebulin interacts with tropomodulin at the pointed ends of the thin filaments. J. Biol. Chem. 276, 583–592 (2001).

16. Pappas, C. T., Bhattacharya, N., Cooper, J. A. & Gregorio, C. C. Nebulin Interacts with CapZ and Regulates Thin Filament Architecture within the Z-Disc. Mol. Biol. Cell 19, 1837–1847 (2008).

17. Labeit, S. et al. Evidence that nebulin is a protein-ruler in muscle thin filaments. FEBS Lett. 282, 313–316 (1991).

18. Gokhin, D. S. et al. Thin-filament length correlates with fiber type in human skeletal muscle. Am. J. Physiol. - Cell Physiol. 302, 555–565 (2012).

19. Kiss, B. et al. Nebulin and Lmod2 are critical for specifying thin-filament length in skeletal muscle. Sci. Adv. 6, 1–18 (2020).

20. Fowler, V. M., McKeown, C. R. & Fischer, R. S. Nebulin: Does It Measure up as a Ruler? Curr. Biol. 16, 18–20 (2006).

21. Bang, M. L. et al. Nebulin-deficient mice exhibit shorter thin filament lengths and reduced contractile function in skeletal muscle. J. Cell Biol. 173, 905–916 (2006).

22. Gonsior, S. M., Gautel, M. & Hinssen, H. A six-module human nebulin fragment bundles actin filaments and induces actin polymerization. J. Muscle Res. Cell Motil. 19, 225–235 (1998).

23. Pfuhl, M., Winder, S. J. & Pastore, A. Nebulin, a helical actin binding protein. EMBO J. 13, 1782–1789 (1994).

24. Pospich, S., Merino, F. & Raunser, S. Structural Effects and Functional Implications of Phalloidin and Jasplakinolide Binding to Actin Filaments. Structure 28, 437-449.e5 (2020).

25. Ao, X. & Lehrer, S. S. Phalloidin unzips nebulin from thin filaments in skeletal myofibrils. J. Cell Sci. 108, 3397–3403 (1995).

26. Castillo, A., Nowak, R., Littlefield, K. P., Fowler, V. M. & Littlefield, R. S. A nebulin ruler does not dictate thin filament lengths. Biophys. J. 96, 1856–1865 (2009).

27. Kruger, M., Wright, J. & Wang, K. Nebulin as a length regulator of thin filaments of vertebrate skeletal muscles: Correlation of thin filament length, nebulin size, and epitope profile. J. Cell Biol. 115, 97–107 (1991).

28. Lukoyanova, N. et al. Each actin subunit has three nebulin binding sites: Implications for steric blocking. Curr. Biol. 12, 383–388 (2002).

29. Poole, K. J. V. et al. A comparison of muscle thin filament models obtained from electron microscopy reconstructions and low-angle X-ray fibre diagrams from non-overlap muscle. J. Struct. Biol. 155, 273–284 (2006).

30. Marttila, M. et al. Nebulin interactions with actin and tropomyosin are altered by disease-causing mutations. Skelet. Muscle 4, 1–10 (2014).

31. Zhang, J. Q., Weisberg, A. & Horowits, R. Expression and purification of large nebulin fragments and their interaction with actin. Biophys. J. 74, 349–359 (1998).

32. Chitose, R. et al. Isolation of nebulin from rabbit skeletal muscle and its interaction with actin. J. Biomed. Biotechnol. 2010, (2010).

33. Jumper, J. et al. Highly accurate protein structure prediction with AlphaFold. Nature (2021) doi:10.1038/s41586-021-03819-2.

34. Manning, E. P., Tardiff, J. C. & Schwartz, S. D. A model of calcium activation of the cardiac thin filament. Biochemistry 50, 7405–7413 (2011).

35. Yamada, Y., Namba, K. & Fujii, T. Cardiac muscle thin filament structures reveal calcium regulatory mechanism. Nat. Commun. 11, 1–3 (2020).

36. Risi, C. M. et al. The structure of the native cardiac thin filament at systolic Ca2+ levels. Proc. Natl. Acad. Sci. U. S. A. 118, 2–9 (2021).

37. Bang, M.-L. et al. Nebulin plays a direct role in promoting strong actin myosin interactions. FASEB J. 23, 4117–4125 (2009).

38. Chandra, M. et al. Nebulin alters cross-bridge cycling kinetics and increases thin filament activation. A novel mechanism for increasing tension and reducing tension cost. J. Biol. Chem. 284, 30889–30896 (2009).

39. Kiss, B. et al. Nebulin stiffens the thin filament and augments cross-bridge interaction in skeletal muscle. Proc. Natl. Acad. Sci. U. S. A. 115, 10369–10374 (2018).

40. Manning, E. P., Tardiff, J. C. & Schwartz, S. D. Molecular effects of familial hypertrophic cardiomyopathy-related mutations in the TNT1 domain of cTnT. J. Mol. Biol. 421, 54–66 (2012).

41. Lehtokari, V. L. et al. Mutation update: The spectra of nebulin variants and associated myopathies. Hum. Mutat. 35, 1418–1426 (2014).

42. Wang, Z. et al. The molecular basis for sarcomere organization in vertebrate skeletal muscle. Cell 184, 2135-2150.e13 (2021).

43. Yang, S. et al. Cryo-EM structure of the inhibited (10S) form of myosin II. Nature 588, 521–525 (2020).

44. Scarff, C. A. et al. Structure of the shutdown state of myosin-2. Nature 588, 515–520 (2020).

## References

45. Solaro, R. J., Pang, D. C. & Briggs, F. N. The purification of cardiac myofibrils with Triton X-100. BBA - Bioenerg. 245, 259–262 (1971).

46. Tacke, S. et al. A streamlined workflow for automated cryo focused ion beam milling. J. Struct. Biol. 107743 (2021) doi:10.1016/j.jsb.2021.107743.

47. Mastronarde, D. N. Automated electron microscope tomography using robust prediction of specimen movements. J. Struct. Biol. 152, 36–51 (2005).

48. Yang, Y. et al. Rigor-like Structures from Muscle Myosins Reveal Key Mechanical Elements in the Transduction Pathways of This Allosteric Motor. Structure 15, 553–564 (2007).

49. Hagen, W. J. H., Wan, W. & Briggs, J. A. G. Implementation of a cryo-electron tomography tilt-scheme optimized for high resolution subtomogram averaging. J. Struct. Biol. 197, 191–198 (2017).

50. Zheng, S. Q. et al. MotionCor2: Anisotropic correction of beam-induced motion for improved cryo-electron microscopy. Nat. Methods 14, 331–332 (2017).

51. Kremer, J. R., Mastronarde, D. N. & McIntosh, J. R. Computer visualization of three-dimensional image data using IMOD. J. Struct. Biol. 116, 71–76 (1996).

52. Tinevez, J. Y. et al. TrackMate: An open and extensible platform for single-particle tracking. Methods 115, 80–90 (2017).

53. Schindelin, J. et al. Fiji: An open-source platform for biological-image analysis. Nat. Methods 9, 676–682 (2012).

54. Schneider, C. A., Rasband, W. S. & Eliceiri, K. W. NIH Image to ImageJ: 25 years of image analysis. Nat. Methods 9, 671–675 (2012).

55. Wagner, T. et al. SPHIRE-crYOLO is a fast and accurate fully automated particle picker for cryo-EM. Commun. Biol. 2, 1–13 (2019).

56. Bharat, T. A. M. & Scheres, S. H. W. Resolving macromolecular structures from electron cryotomography data using subtomogram averaging in RELION. Nat. Protoc. 11, 2054–2065 (2016).

57. Yang, Z., Fang, J., Chittuluru, J., Asturias, F. J. & Penczek, P. A. Iterative stable alignment and clustering of 2D transmission electron microscope images. Structure 20, 237–247 (2012).

58. Tegunov, D., Xue, L., Dienemann, C., Cramer, P. & Mahamid, J. Multi-particle cryo-EM refinement with M visualizes ribosome-antibiotic complex at 3.5 Å in cells. Nat. Methods 18, 186–193 (2021).

59. Tegunov, D. & Cramer, P. Real-time cryo-electron microscopy data preprocessing with Warp. Nat. Methods 16, 1146–1152 (2019).

60. Moriya, T. et al. High-resolution single particle analysis from electron cryo-microscopy images using SPHIRE. J. Vis. Exp. 2017, 1–11 (2017).

61. Terwilliger, T. C., Ludtke, S. J., Read, R. J., Adams, P. D. & Afonine, P. V. Improvement of cryo-EM maps by density modification. Nat. Methods 17, 923–927 (2020).

62. Eswar, N., Eramian, D., Webb, B., Shen, M.-Y. & Sali, A. Protein Structure Modeling with MODELLER. in Structural Proteomics: High-Throughput Methods (eds. Kobe, B., Guss, M. & Huber, T.) 145–159 (Humana Press, 2008). doi:10.1007/978-1-60327-058-8_8.

63. von der Ecken, J., Heissler, S. M., Pathan-Chhatbar, S., Manstein, D. J. & Raunser, S. Cryo-EM structure of a human cytoplasmic actomyosin complex at near-atomic resolution. Nature 534, 724–728 (2016).

64. Emsley, P., Lohkamp, B., Scott, W. G. & Cowtan, K. Features and development of Coot. Acta Crystallogr. Sect. D Biol. Crystallogr. 66, 486–501 (2010).

65. Croll, T. I. {\it ISOLDE}: a physically realistic environment for model building into low-resolution electron-density maps. Acta Crystallogr. Sect. D 74, 519–530 (2018).

66. Goddard, T. D. et al. UCSF ChimeraX: Meeting modern challenges in visualization and analysis. Protein Sci. 27, 14–25 (2018).

67. Afonine, P. V et al. New tools for the analysis and validation of cryo-EM maps and atomic models. Acta Crystallogr. Sect. D 74, 814–840 (2018).

68. Chen, V. B. et al. {\it MolProbity}: all-atom structure validation for macromolecular crystallography. Acta Crystallogr. Sect. D 66, 12–21 (2010).

69. Barad, B. A. et al. EMRinger: Side chain-directed model and map validation for 3D cryo-electron microscopy. Nat. Methods 12, 943–946 (2015).

70. Waterhouse, A. et al. SWISS-MODEL: Homology modelling of protein structures and complexes. Nucleic Acids Res. 46, W296–W303 (2018).

71. Larkin, M. A. et al. Clustal W and Clustal X version 2.0. Bioinformatics 23, 2947–2948 (2007).

72. Crooks, G. E., Hon, G., Chandonia, J. M. & Brenner, S. E. WebLogo: A sequence logo generator. Genome Res. 14, 1188–1190 (2004).

73. Wang, S., Li, W., Liu, S. & Xu, J. RaptorX-Property: a web server for protein structure property prediction. Nucleic Acids Res. 44, W430–W435 (2016).

74. Stan Development Team. Stan Modeling Language Users Guide and Reference Manual, 2.27 (2019). https://mc-stan.org

75. Gasteiger, E. et al. Protein Identification and Analysis Tools on the ExPASy Server. in The Proteomics Protocols Handbook (ed. Walker, J. M.) 571–607 (Humana Press, 2005). doi:10.1385/1-59259-890-0:571.

76. Abraham, D. J. & Leo, A. J. Extension of the fragment method to calculate amino acid zwitterion and side chain partition coefficients. Proteins Struct. Funct. Bioinforma. 2, 130–152 (1987).

